# Bayesian modeling of BAC firing as a mechanism for apical amplification in neocortical pyramidal neurons

**DOI:** 10.1101/604066

**Authors:** Jim W. Kay, W. A. Phillips, Jaan Aru, Bruce P. Graham, Matthew E. Larkum

## Abstract

Pyramidal cells in layer 5 of the neocortex have two distinct integration sites. These cells integrate inputs to basal dendrites in the soma while integrating inputs to the tuft in a site at the top of the apical trunk. The two sites communicate by action potentials that backpropagate to the apical site and by backpropagation-activated calcium spikes (BAC firing) that travel from the apical to the somatic site. Six key messages arise from the probabilistic information-theoretic analyses of BAC firing presented here. First, we suggest that pyramidal neurons with BAC firing could convert the odds in favour of the presence of a feature given the basal data into the odds in favour of the presence of a feature given the basal data and the apical input, by a simple Bayesian calculation. Second, the strength of the cell’s response to basal input can be amplified when relevant to the current context, as specified by the apical input, without corrupting the message that it sends. Third, these analyses show rigorously how this apical amplification depends upon communication between the sites. Fourth, we use data on action potentials from a very detailed multi-compartmental biophysical model to study our general model in a more realistic setting, and demonstrate that it describes the data well. Fifth, this form of BAC firing meets criteria for distinguishing modulatory from driving interactions that have been specified using recent definitions of multivariate mutual information. Sixth, our general decomposition can be extended to cases where, instead of being purely driving or purely amplifying, apical and basal inputs can be partly driving and partly amplifying to various extents. These conclusions imply that an advance beyond the assumption of a single site of integration within pyramidal cells is needed, and suggest that the evolutionary success of neocortex may depend upon the cellular mechanisms of context-sensitive selective amplification hypothesized here.

**Author summary:** The cerebral cortex has a key role in conscious perception, thought, and action, and is predominantly composed of a particular kind of neuron: the pyramidal cells. The distinct shape of the pyramidal neuron with a long dendritic shaft separating two regions of profuse dendrites allows them to integrate inputs to the two regions separately and combine the results non-linearly to produce output. Here we show how inputs to this more distant site strengthen the cell’s output when it is relevant to the current task and environment. By showing that such neurons have capabilities that transcend those of neurons with the single site of integration assumed by many neuroscientists, this ‘splitting of the neuronal atom’ offers a radically new viewpoint from which to understand the evolution of the cortex and some of its many pathologies. This also suggests that approaches to artificial intelligence using neural networks might come closer to something analogous to real intelligence, if, instead of basing them on processing elements with a single site of integration, they were based on elements with two sites, as in cortex.

## Introduction

Contextual disambiguation is crucial to cognition in general and perception in particular [1], as highlighted by the work of many artists such as René Magritte and Maurits Escher. It can be considered to be a context-sensitive form of Bayesian inference because it uses prior knowledge to make inferences from new data. Context-sensitive selection applies not only to conscious experience and behavior but also to individual neurons [2, 3]. A plausible neuronal mechanism for context-sensitive selective amplification in cortical pyramidal neurons that exploits the architecture of long-range connectivity in the cortex and the intrinsic properties of some pyramidal cell dendrites has been proposed [4]. Contextual information arriving at the neuron’s apical tuft can activate the apical integration site (AIS) that generates long-lasting dendritic spikes. This process, however, is contingent on the presence of back-propagating action potentials that lower the threshold in the AIS (‘BAC firing’ [4, 5, 6, 7, 9, 10]). Thus, as argued elsewhere [11], the two-point neurons hypothesized here challenge the assumption held by many [12] that, from a system point of view, neurons in general operate as integrate-and-fire processors with a single point of integration.

According to our hypotheses, ‘contextual disambiguation’ cannot be identified with a particular kind of information, such as information about the particular time and place at which events occur. It is a particular way of using information. Contextual modulation in two-point neurons influences the AIS to amplify (or attenuate) transmission of information about the cell’s basal and perisomatic inputs. Perisomatic inputs are assumed to convey information about data specific to the features being processed in that region (or column) of the cortex. From this perspective the notion of ambiguity can be interpreted broadly to include ambiguities of a signal’s presence, identity, emotional valence, and relevance to current goals.

A currently prominent view of neocortical function suggests that context-sensitive integration of prior knowledge and sensory data occurs in a Bayesian fashion. Here, we show how the function of BAC firing can be interpreted in context-sensitive Bayesian terms because it is contingent on internal knowledge stored in the strengths of the basal and apical synapses. In brief, we suggest that the output of a pyramidal neuron can be interpreted as approximating the probability of a (columnar) feature given both the basal input and the context and that it is facilitated by the apical dendrite that approximately computes the probability of the context given the feature and the basal input. Unlike two neurons connected via synapses, the two dendritic compartments are linked via a privileged connection, the apical trunk, that allows non-linear bi-directional signalling via active dendritic currents and spikes. This therefore endows the pyramidal neuron with processing capabilities for context-selective amplification that would otherwise require more sophisticated neural circuitry and thus more time and energy.

The hypotheses presented here assume that learning can be viewed as statistical representation of feature-related and contextual information. It has previously been shown that, given functionally distinct receptive and contextual field inputs, learning rules can change synaptic strengths so as to adapt them to the latent statistical structure in their inputs [15, 13, 14, 16]. It has been suggested that BAC firing can provide the activation functions used in those theories [17], but till now that has not been shown explicitly. Furthermore, though the pyramidal cell as a whole can be analyzed as an activation function with two inputs and one output, we show here that it can be more explicitly related to BAC firing by describing each of the somatic and apical integration sites as operating as a three-term activation function with one input from outside the cell and one input from inside the cell. This more realistic representation may not be as complicated as it sounds, however, because we show that the somatic site, the apical site and the cell as a whole can all be described using the same general form of activation function. Though BAC firing has previously been studied using computational models [7, 10], it has not till now been explicitly related to Bayesian inference.

It is known that single spikes are unreliable and having several spikes within a short window (a burst) increases the reliability of information transfer and thus bursts may make a special contribution to brain function [18]. Hence, the fact that apical activity greatly enhances the probability of a second spike makes the output of the cell more reliable and more likely to contribute to perception [19] and behavior [20]. To further simplify the analyses we build on the fact that the probabilities being estimated are of a binary event, i.e. whether or not a second AP is generated. This enables us to use the ratio of these two probabilities, known as the odds, and the log of that ratio, known as the log odds. Though less intuitive, the log odds provides a simple description of a wide range of physiological and psychophysical phenomena [21]. In our application, the basal input is taken into account before the apical input so, using Bayes’ theorem, the posterior log odds given both these inputs can be written as a sum of three terms: the prior log odds, the weight of evidence in favor of a second AP provided by the basal input, and the additional weight of evidence in favor of a second AP provided by the apical input given the basal input. The weight of evidence was first used in [22]. Such expressions of the posterior odds are found in [23, 25, 24], where their use is attributed to A. M. Turing in the 1940s; see also [26].

Thus, our approach contradicts four widely held assumptions. First, we deny that neurons in general function as point processors. Second, we deny that Bayesian-like computations necessarily imply supra-neuronal generative systems. Third, we deny that if a Bayesian inference is sensitive to context then the terms in the Bayesian inference must all be conditioned on context, as proposed by Lee and Mumford [27]. Instead, we argue that biological plausibility and functional potential are enhanced by first conditioning on the basal input. Fourth, we deny that amplification can be identified with multiplicative interactions, which are symmetric, and which have been found to transmit no information unique to either of the interacting terms [28]. In contrast to that, the apical amplification that we hypothesize is asymmetric, and it has been found [29] to transmit information unique to the basal but not to the apical input. This last point is crucial to our approach because it shows how a neuron’s output can be amplified or attenuated by variables that do not thereby become part of the essential information content of the neuron’s output.

The main body of the paper is organized as follows. We begin by presenting a simple idealized sketch of inputs to and outputs from the two integration sites and of the interaction between them via the apical trunk. A Bayesian interpretation of the intra-cellular computation is then introduced and decompositions of log odds are presented. General activation functions are introduced and then used in a comparison of two models for the analysis of binarised data from a detailed multi-compartment model of apical function developed in [10]. A categorised version of the data from [10] is then analysed using information theory and partial information decomposition [30]. Finally, results are presented for general decompositions of log odds for cases in which the basal and apical inputs may be partly driving and partly amplifying to various extents.

## Results

### Two sites of integration that interact via backpropagation and BAC firing in layer 5 pyramidal cells

An idealization of integration within and communication between the somatic and apical sites is shown in Fig 1. This is designed to relate known physiological processes to the probabilistic analyses presented in the following sections. Each integration site receives input from two sources, one that is extracellular and one that is intracellular. The somatic site receives extracellular input from basal/perisomatic synapses, and we refer to it as the ‘basal input’, *b*. It also receives internal input, *c*, from the apical site. The apical site receives extracellular input from apical synapses in layer 1, which we refer to as the ‘apical input’, *a*. It also receives internal input, *z*_1_, in the form of the action potential that is backpropagated from the somatic site.

**Figure 1:**
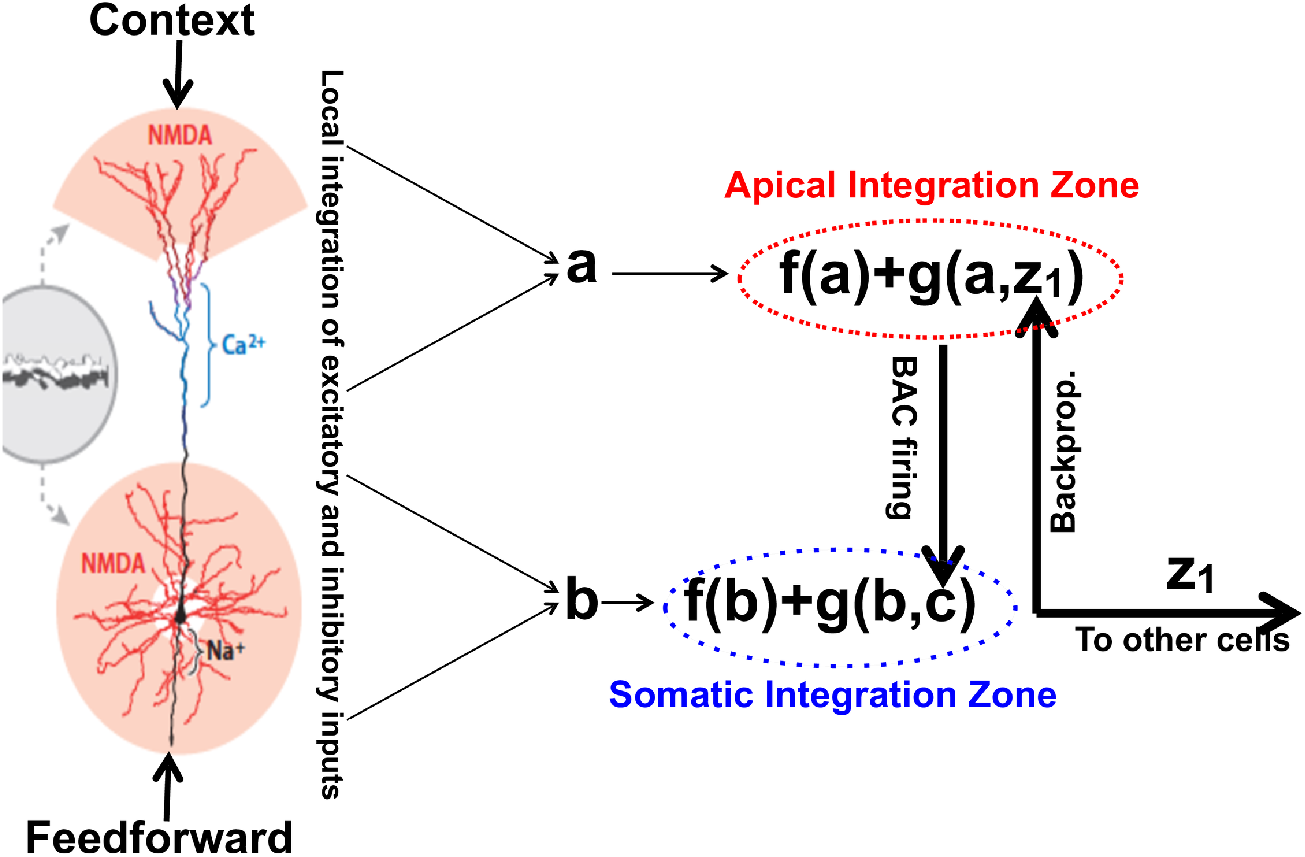
Integration within and between somatic and apical sites. An idealization of integration within and communication between somatic and apical sites in the case where an initiating bAP has been generated and BAC firing is triggered. *z*_1_ is a binary variable that equals 1 if there has been a somatic action potential and zero if not. A backpropagated action potential amplifies transmission of information, via *c* = *f* (*a*) + *g*(*a, z*_1_), about the apical input, *a*, to the somatic integration site, where it is used to amplify transmission of information about the basal input, *b*, to other cells. The functions, *f, g*, are defined below. (The layer 5 pyramidal cell on the left is from Fig 4G of [31].)

Basal dendrites typically receive their input from a few narrowly specified feedforward sources that in sensory and perceptual regions ascend the hierarchy of neocortical abstraction. Apical dendrites typically receive input from diverse sources that include feedback from higher neocortical regions, higher-order thalamus, the amygdala, and the adrenergic and cholinergic systems. This diverse set of sources provides contextual information that guides processing and learning of the feedforward data. For simplicity we refer to this set of diverse inputs collectively as ‘context’. It amplifies response to feedforward signals that are relevant given that particular context. We refer to the post-synaptic locations that receive feedforward information as ‘basal’, however they may also include perisomatic locations, as shown in Fig 1. Positive values of *a* and *b* are taken to be analogous to net depolarization at that site, and negative values to hyperpolarization. Computation of the net basal and apical inputs are shown in Fig 1 as occurring outside their respective sites of integration, but we assume that it occurs as part of the integration within sites.

Empirical data indicates that BAC firing most frequently adds one more AP to the initiating AP; sometimes two; and rarely more. We assume that our focus on the 10 ms time-scale is justified because two or more spikes within about 10 ms is a faster and more energy efficient signal than mean spike rate over 100 ms, and because both data and models indicate that apical depolarization has more effect on bursting than on mean spike rate (e.g. [10,8,9]). To extend our analysis to adequately cover bursts of different lengths, inter-burst intervals, and fast regular spiking we would have to include the effects of negative feedback from inhibitory interneurons, including those specific to the tuft. The refractory nature of AP generation and synaptic facilitation/accomodation would also become relevant if we were to present the analysis as being concerned with a sequence of APs. The present analysis makes no assumptions about those things but simply considers the dependence of AP probability on *a* and b given that a back-propagating action potential (bAP) has occurred. There are subtle simplifications in the idealization presented, e.g. *a* and *b* are considered to be stable on the brief time-course of the BAC firing effect. We first discuss the case where the apical input provides amplifying ‘context’, while the basal input is driving.

### A Bayesian interpretation of intra-site computation

We represent the net apical input by the continuous random variable, *A*, which is a weighted and summed combination of inputs from various sources in the apical dendrites. The observed value of *A* is the *a* used in Fig 1. We represent the net basal input by the continuous random variable, *B*, which is a weighted and summed combination of inputs from various sources in the basal dendrites. The observed value of *B* is the *b* used in Fig 1. The binary random variable *Z*_1_ denotes whether or not a first, initiating bAP is generated; its observed value, *z*_1_, is used in Fig 1, and *z*_1_ = 1 when an initiating bAP has been generated. *Z*_2_ indicates whether or not a second AP is emitted. When discussing odds and log odds in the sequel the symbol, *S*_2_, will denote the event *Z*_2_ = 1, with 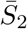 as the complementary event. We use the symbol ‘*P*’ for probabilities, given the discrete nature of the *Z_i_*, and ‘*p*’ for continuous probability density functions.

We consider first the scenario in which the BAC firing in the apical site has been triggered by the arrival of an initiating bAP, as illustrated in Fig 1. We then focus on the resulting computation in the somatic site. We have a prior distribution on *Z*_2_, with *O*(*S*_2_) denoting the prior odds in favour of a second AP, and a generative model for the basal input, *b*, given that *Z*_2_ = *z*_2_, where *z*_2_ = 0 or 1. These are combined using Bayes’ theorem to produce the posterior probability that *Z*_2_ = 1 given the basal input, *b*:

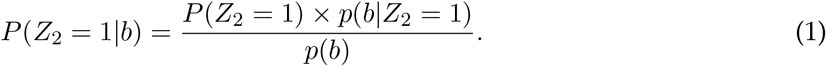

As explained in the Methods section, we can use (1) to deduce the following relationships between prior and posterior odds and log odds in favour of a second AP.

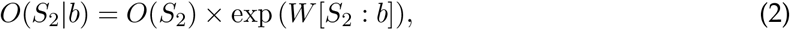

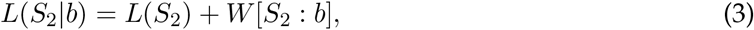

where *W* [*S*_2_: *b*] is the weight of evidence in favour of a second AP that is provided by the basal input.

Then the posterior probability in (1) is updated by taking into account the observed apical input, *a*, and using Bayes’ theorem to give

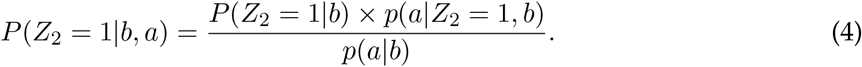

In (4), *p*(*a|Z*_2_ = 1, *b*) is a generative model for the apical input given the observed basal input, *b*, and the event *Z*_2_ = 1. Using (4), we can write the updated odds and log odds as follows.

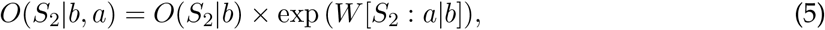

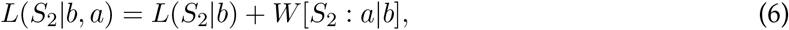

where *W* [*S*_2_: *a|b*] is the weight of evidence in favour a second AP that is provided by the apical input, given the basal input.

Using (2), (5), we offer a Bayesian interpretation of computation within the somatic site, once BAC firing has been triggered. We start with the prior odds in favour of a second AP, *O*(*S*_2_). Taking the basal input into account and using Bayes’ theorem, this prior odds is updated to form the posterior odds in favour of a second AP, *O*(*S*_2_|*b*), given the basal input. These posterior odds will increase (decrease) when the weight of evidence in favour of a second AP provided by the basal input is positive (negative). Then the posterior odds *O*(*S*_2_|*b*) is updated by computing the posterior odds in favour of a second AP, *O*(*S*_2_|*b, a*), given also the apical input. The odds *O*(*S*_2_|*b*) are thus amplified or attenuated depending on whether the weight of evidence provided by the apical input in addition to the basal input is positive or negative.

From (3), (6) we can summarize this Bayesian interpretation as a three-term additive decomposition of the posterior log odds in favour of a second AP,

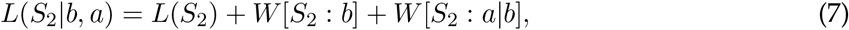

in terms of the prior log odds in favour of a second AP, the weight of evidence in favour of a second AP provided by the basal input and the weight of evidence in favour of a second AP provided by the apical input in addition to the basal input. This computation relates to the somatic integration site, but the term *W* [*S: a*| *b*] would not be available without the two-way communication between the sites by which the increased apical activation is transmitted to the somatic site, thus increasing the odds of the propagation of a second AP within a time interval of around 10 ms. An illustration of the implementation of the Bayesian interpretation is provided in S2 File.

### General decomposition of log odds

In Eqs (2), (7), based on the specification of generative models for the basal and apical inputs, the weight of evidence term *W* [*S*_2_: *b*] is a function of *b* and *W* [*S*_2_: *a: b*] is a function of both *a* and *b*. By analogy, and not based on generative models, we now define expressions of the log odds in terms of general activation functions *f* (*b*) and *g*(*a, b*) that will be used in the sequel.

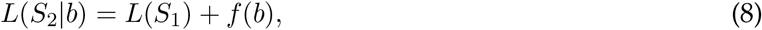

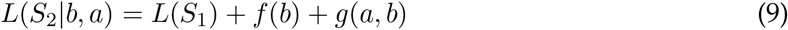

The general functions employed have the following forms.

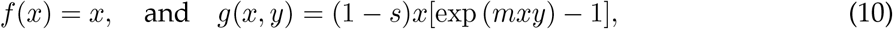

where 0 < *s* < 1 and *m* > 0, and we set the free constants, *s, m*, using the values 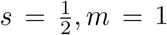, although in the work described below in the Bayesian Modeling subsection it is necessary to employ much smaller values for *m*. The specific forms of activation function used are given in Table 1, and further discussion is provided in the Methods section.

**Table 1:** Specific activation functions. Specific expressions for the activation functions in the apical and somatic sites during Phase 1 (when no initiating action potential is emitted from the basal site) and Phase 2 (when an initiating bAP has been received from the basal site and BAC firing has been triggered)

|  | Phase 1 | Phase 2 |
| --- | --- | --- |
| Site | $z_1 = 0$ | $z_1 = 1$ |
| Apical | $a$ | $\frac{1}{2}a(1 + e^a)$ |
| Somatic | $b$ | $\frac{1}{2}b[1 + \exp[\frac{1}{2}ba(1 + e^a)]]$ |

In the following subsections, we discuss computation in the somatic integration site (SIS) and make use of activation functions in which the apical input is purely amplifying and the basal input is purely driving. In the final subsection on alternative modes of apical function, we consider activation functions in which the basal and apical inputs may be partly driving and partly amplifying.

### Dependence of second AP probability on apical and basal input

We now consider general expressions for the posterior probabilities of a second AP, given basal input, and given both basal and apical input, respectively. First we note that *P*(*S*_2_) gives the prior probability of a second AP, which we set to 0.005.

Therefore, we may write

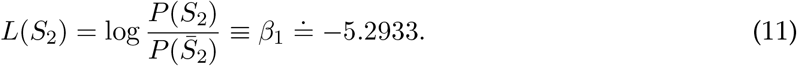

Using, from Table 1,

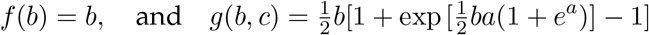

in Eqs (8), (9) we have that

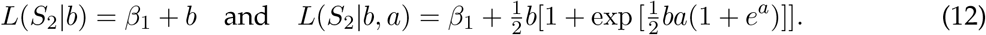

By using the connection between probability and log odds, we can write the posterior probabilities as

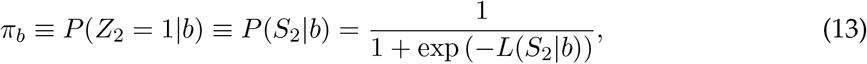

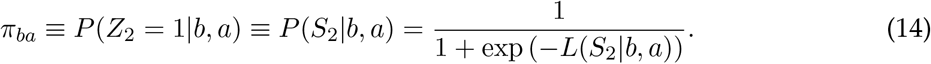

The posterior probabilities *π_b_*, given only the basal input, and *π_ba_*, given both basal and apical inputs, are shown for different positive strengths of apical input in Fig 2.

**Figure 2:**
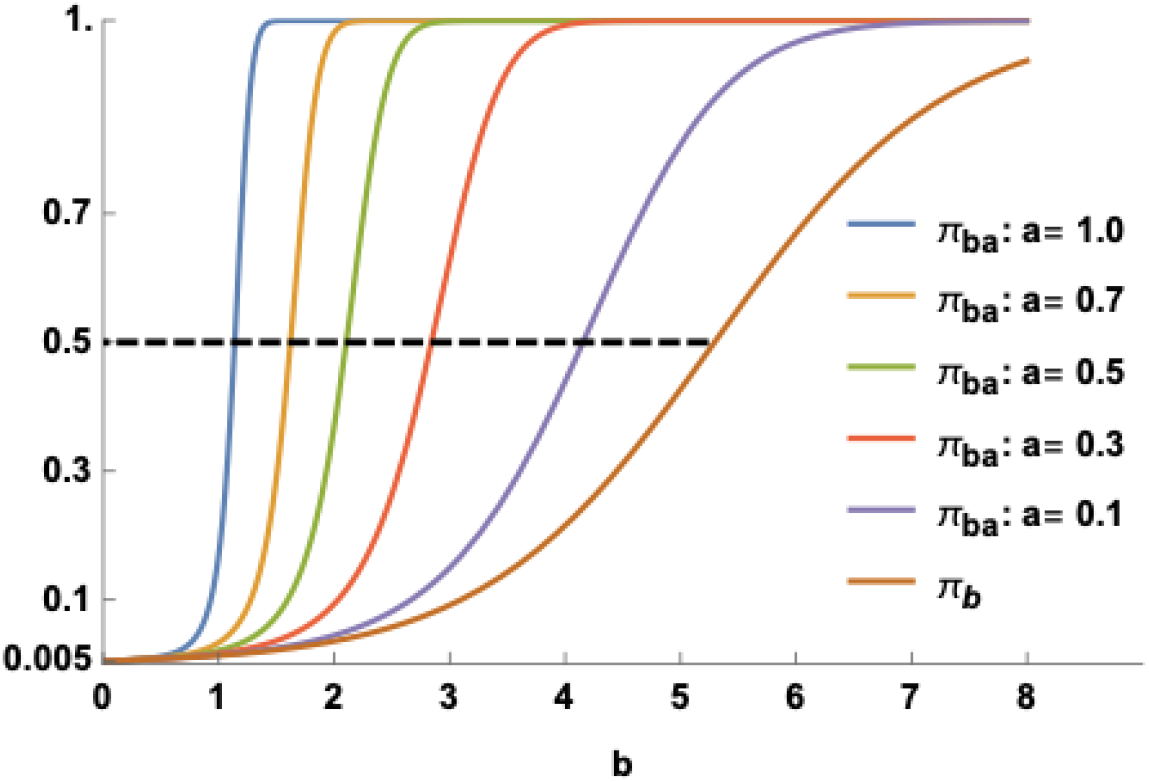
Dependence of second AP probability on apical and basal input. Posterior probabilities, *π_a_*, *π_b_*, are plotted against positive values of the basal input, *b*, using five different values of the apical input, *a*. The dashed line indicates points on the curves at which the posterior probability is 0.5. The typical baseline firing probability, *P*(*S*_2_), is assumed to be 0.005.

We see that due to apical amplification the posterior probability, *π_ba_*, of a second AP given both apical and basal input is larger than the corresponding posterior probability, *π_b_*, given only the basal input. Apical input has little or no effect when basal input is very weak or very strong, but has a dramatic effect at intermediate values of basal input strength. This supralinear effect becomes more pronounced as the strength of the apical input increases. Defining the basal threshold as that value of the basal input for which the posterior probability is 0.5, we notice, in particular, that this threshold decreases markedly as the strength of the apical input increases. This effect of apical input on response to basal input is well matched to that sketched in Fig 2c of [4]. It is also well matched to the performance of a two ‘compartment’ model (e.g. Figure 5b of [32]) which shows that apical amplification depends strongly on calcium conductance in the apical trunk.

### Bayesian modelling of the binarised action potential data from a detailed multi-compartment model

Shai et al. [10] used a multi-compartmental model to produce data on the frequency of somatic spike output for 31 given numbers of basal inputs, 0,10,20,…, 300, equally spaced between 0 and 300, and 21 given numbers of apical tuft inputs, 0,10,20,…, 200, equally spaced between 0 and 200. They then developed a phenomenological composite sigmoidal model for the frequency data, thus providing an explanation for coincidence detection between basal and apical tufts. They also produced data on the number of action potentials for the same given combinations of basal and apical tuft inputs, but they did not report any analysis of these data.

Our interest lies in modeling the posterior probability of a second AP within around 10 ms since an initial bAP has been generated. The number of APs in the data file ranges from 0 to 4. We are interested in the occurrence of a second AP, i.e. 2 APs, and this event happens also when 3 or 4 APs are observed. The data were therefore recoded by setting 0 or 1 AP to 0 and 2-4 APs to 1, thus creating a binary matrix where a ‘1’ denotes the occurrence of a second AP and ‘0’ means that this event has not happened. An alternative interpretation of these binary data, for each combination of the numbers of basal and apical inputs, is that a ‘1’ indicates that bursting (2-4 APs) has happened while a ‘0’ means ‘no bursting’ (0-1 APs).

The data are shown in Fig 3A. For each number of tuft inputs, the points of transition from ‘blue’ to ‘red’ along each row give a noisy indication of the threshold – the value of the basal input for which the posterior probability is equal to 0.5. It is clear from the plot that the thresholds vary according to the number of tuft inputs: the threshold is large when the number of tuft inputs is low whereas it is low when the number of tuft inputs is large.

**Figure 3:**
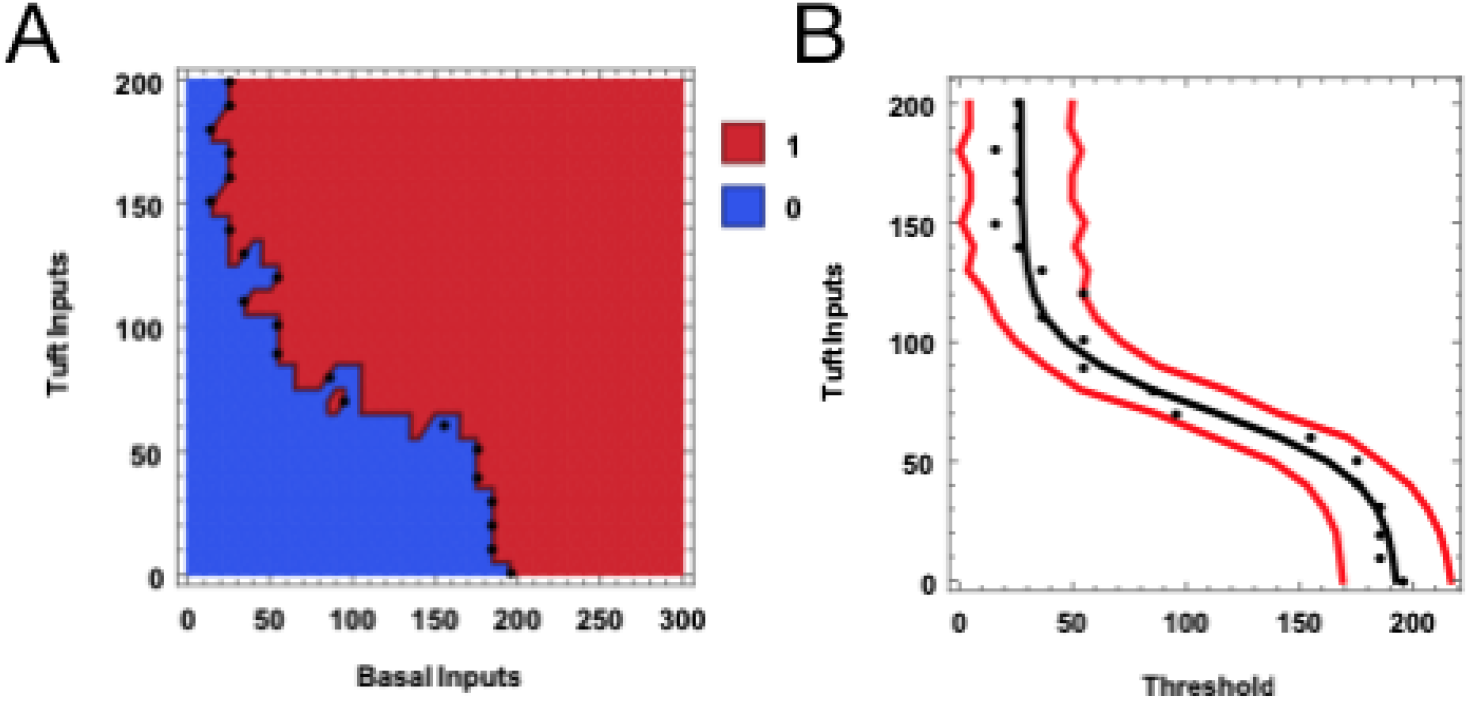
The binary data and estimated threshold curve. Illustrations of the binary AP data based on [10]. (A) An image of the binary data on a 21 by 31 grid, with points showing the estimated thresholds. (B) The predicted threshold curve given (in black) as the median of the posterior predictive threshold distribution, together with pointwise 95% posterior predictive intervals (in red) for each number of tuft inputs, obtained by applying weighted Bayesian nonlinear regression with the estimated thresholds (given as points) as responses, the number of tuft inputs as the explanatory variable, and the mean threshold given as a four parameter logistic function of the number of tufts inputs. These intervals were computed for each of the 21 numbers of apical inputs and interpolation used.

We use Bayesian modelling to fit two phenomenological models to these binary data [33] using `rstan` [34]. We take the prior log odds *L*(*S*_2_) to be −5.2993, as before.

### A threshold model

The composite model in [10] is based on two logistic functions *M, T* of the tuft inputs. *M* modeled the maximum AP frequencies for each number of tuft inputs, while *T* modeled the thresholds. Since we consider binary output, for which the maxima are 1 for each number of tuft inputs, there is no need for *M* here (it is 1). We therefore model only the thresholds.

For each of the *n_a_* = 21 apical inputs, *a* = {*a_i_*}, used in [10], a penalized binary logistic regression model was fitted, using ‘R’ [35,36,37,38]. The explanatory variable was the number of basal inputs, {*b_i_*}. The level of basal input for which the probability of a second AP is equal to 0.5 was estimated to give the estimated thresholds, **t** = {*t_i_*}, and the estimated thresholds are shown as points in Figs 3A, B. The estimated standard errors associated with these estimates were noted and used to define weights in the subsequent analysis. The weight, *w_i_*, for each threshold estimate was taken to be the estimated standard error associated with the estimated threshold, *t_i_*, and is assumed to be known. The basal and threshold values were each scaled to lie in the interval [0, 2], and the weights were changed accordingly. A weighted Bayesian nonlinear regression model, with *p*(*t_i_*|*a_i_*, ***θ***) assumed to be a Gaussian pdf with mean

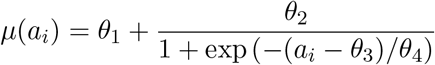

and variance (*σ/w_i_*)^2^, where ***θ*** = [*θ*_1_,*θ*_2_,*θ*_3_,*θ*_4_, *σ*^2^} are unknown parameters. It was assumed that the {t_i_} are conditionally independent given the {*a_i_*} and ***θ***. The *θ_i_* and *σ* parameters were assumed *a priori* to be mutually independent with weakly informative priors: Gaussian with mean 0 and variance 10 for each of the *θ_i_* and Uniform on [0, 20] for *σ*. The posterior distribution for ***θ*** given the data 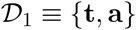 is

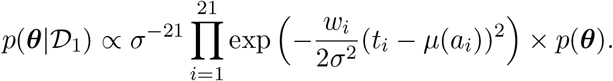

The constraints, *θ*_2_ > 0, *θ*_4_ < 0, were added to ensure that an increasing logistic function was fitted. This model was fitted using `rstan` [34], and large samples from the posterior distributions of the parameters were produced for subsequent computation of the posterior predictive probabilities of a second AP, as described below in Eqs (15) – (18). Further detail is referenced in the supplementary S1 File. Some details of the fitted model are given in Table 2.

**Table 2:**
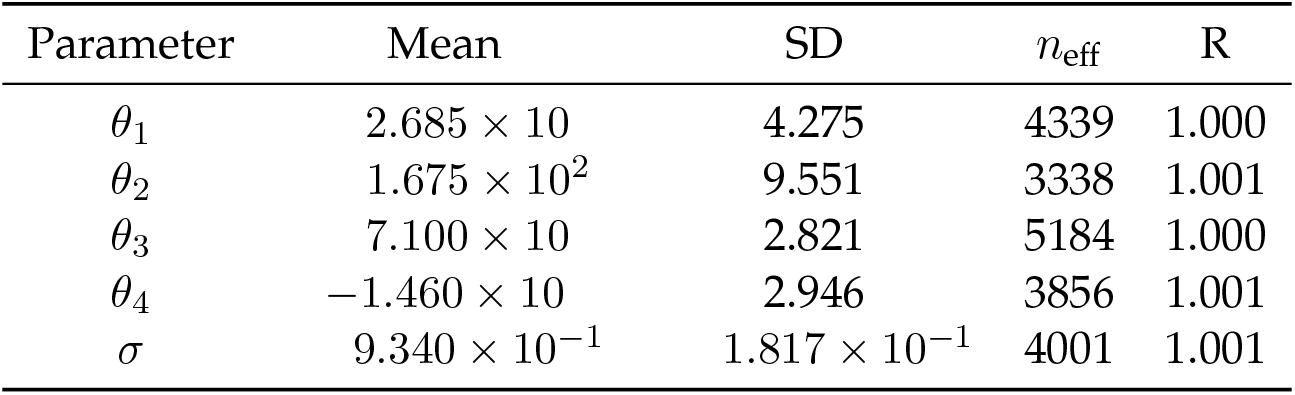
Posterior summary: threshold curve fitting. The posterior mean and standard deviation, based on 10,000 MCMC samples, for each parameter in the nonlinear regression model. Also shown are the effective samples sizes and the values of the statistic *R*, which should be close to 1 if the four chains used have all converged to the same distribution.

It is also of interest to predict the threshold, *t*_new_, given a new value, *a*_new_, of the apical input. The posterior predictive distribution for *t*_new_ is given by

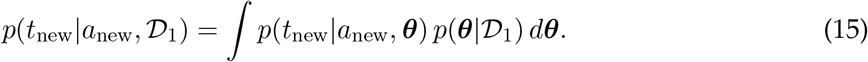

The same Gaussian model, as described above, was assumed for each *t*_new_, given *a*_new_ and ***θ***. For each new apical input, *a*_new_, a large number of values for the predicted threshold were then produced to provide a sampled version of the posterior predictive distribution in Eq 15. The prediction, 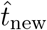, is taken to be the median of these sampled predictions, and pointwise 95% posterior predictive limits were obtained by using the 0.025th and 0.975th quantiles.

The threshold logistic curve given by the median of each posterior predictive threshold distribution is shown in Fig 3B, together with 95% pointwise prediction intervals. The threshold logistic curve decreases monotonically as the number of tuft inputs increases. The pointwise posterior prediction intervals give an indication of the uncertainty of the predicted thresholds. There is greater uncertainty when the number of tuft inputs is larger than 120 or less than 50.

Attention turns now to computing posterior predictive probabilities of a second AP for new values, *a, b,* of the apical and basal input, respectively, given the data, 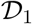. By analogy with the composite model in [10], we define a general decomposition for the log odds in favor of a second AP provided by the net basal and apical inputs in terms of the parameter vector ***θ*** as

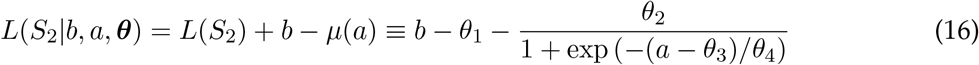

and so the posterior probability of a second AP under this threshold model is

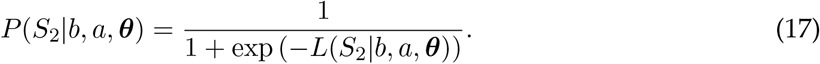

The posterior predictive probability of a second AP for new values *b, a* of basal and apical input, given the data 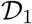 is then

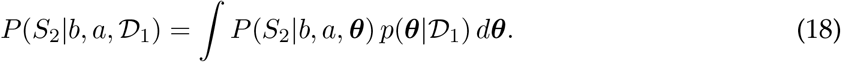

The computation of these probabilities is described in the Methods section.

### A general model

In Fig 3A, we see that there is almost complete separation between the 0’s and the 1’s. Therefore it is necessary to employ regularization, and here we take a Bayesian approach by defining a prior on *β* which enforces this. There are 651 combinations of basal and apical inputs and binary outputs, 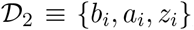, and for each combination there is a binary response, *Z_i_*, that takes the value 1, with probability *p_i_*, when a second AP occurs and 0 otherwise. For each *i, Z_i_* has a Bernoulli distribution with parameter *p_i_*. The {*z_i_*} were assumed to be conditionally independent given the {*a_i_, b_i_*} and ***β***. The binary logistic nonlinear regression model has the form

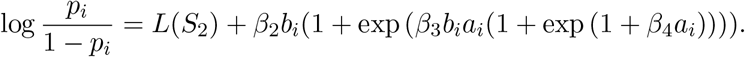

The posterior distribution of ***β*** given the data 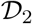 is

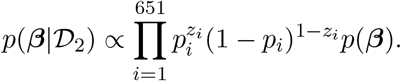

The basal and apical data were scaled to lie in the intervals [0, 2] and [0, 3], respectively. The parameters, *β*_2_, *β*_3_,*β*_4_ were assumed *a priori* to be mutually independent with each following a uniform probability model on (0, 10), which provides a weakly informative prior. This interval is chosen so that the *β_i_* parameters will not be allowed to become too large, which they would otherwise do since there is almost complete separation between the 0’s and the 1’s in the data.

Therefore, the posterior log odds in favor of a second AP under the general model is

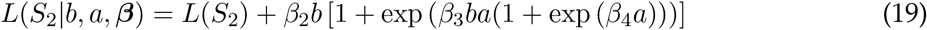

and so the posterior probability of a second AP under this general model is

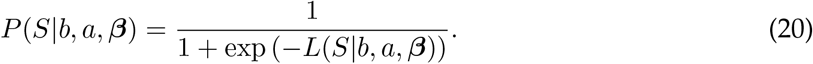

The posterior predictive probability of a second AP for new values, *b, a*, of basal and apical input, given the data 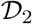 is then obtained by

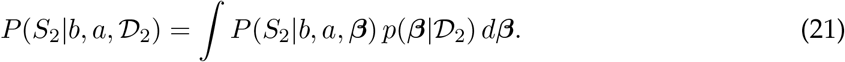

The computation of these probabilities is described in the Methods section. Some details of the fitted model are given in Table 3.

**Table 3:** Posterior summary: General Model. The posterior mean and standard deviation, based on 10,000 MCMC samples, for each parameter in the general model. Also shown are the effective samples sizes and the statistic *R*, which should be close to 1 if the four chains used have all converged to the same distribution.

| Parameter | Mean | SD | $n_{\text{eff}}$ | R |
| --- | --- | --- | --- | --- |
| $\beta_2$ | $1.221 \times 10^{-2}$ | $7.568 \times 10^{-4}$ | 3579 | 1.001 |
| $\beta_3$ | $3.747 \times 10^{-5}$ | $4.860 \times 10^{-6}$ | 2942 | 1.002 |
| $\beta_4$ | $1.565 \times 10^{-2}$ | $7.740 \times 10^{-2}$ | 3238 | 1.001 |

### A comparison of the two models

Contour plots of the posterior predictive probability functions are displayed in Figs 4A, 4B. The regions of very high probability (> 0.9), or very low probability (< 0.1), are not identical but they share similar combinations of numbers of basal and apical tuft inputs, and they have a large overlap; of course, the specialized threshold model gives a better fit for the nonlinear boundary between 0s and 1s. Maximum probability is attained not only when both the basal and apical inputs are large, which indicates a form of coincidence detection, but also when the basal input is large (200-300) while the apical input is low (0-50).

**Figure 4:**
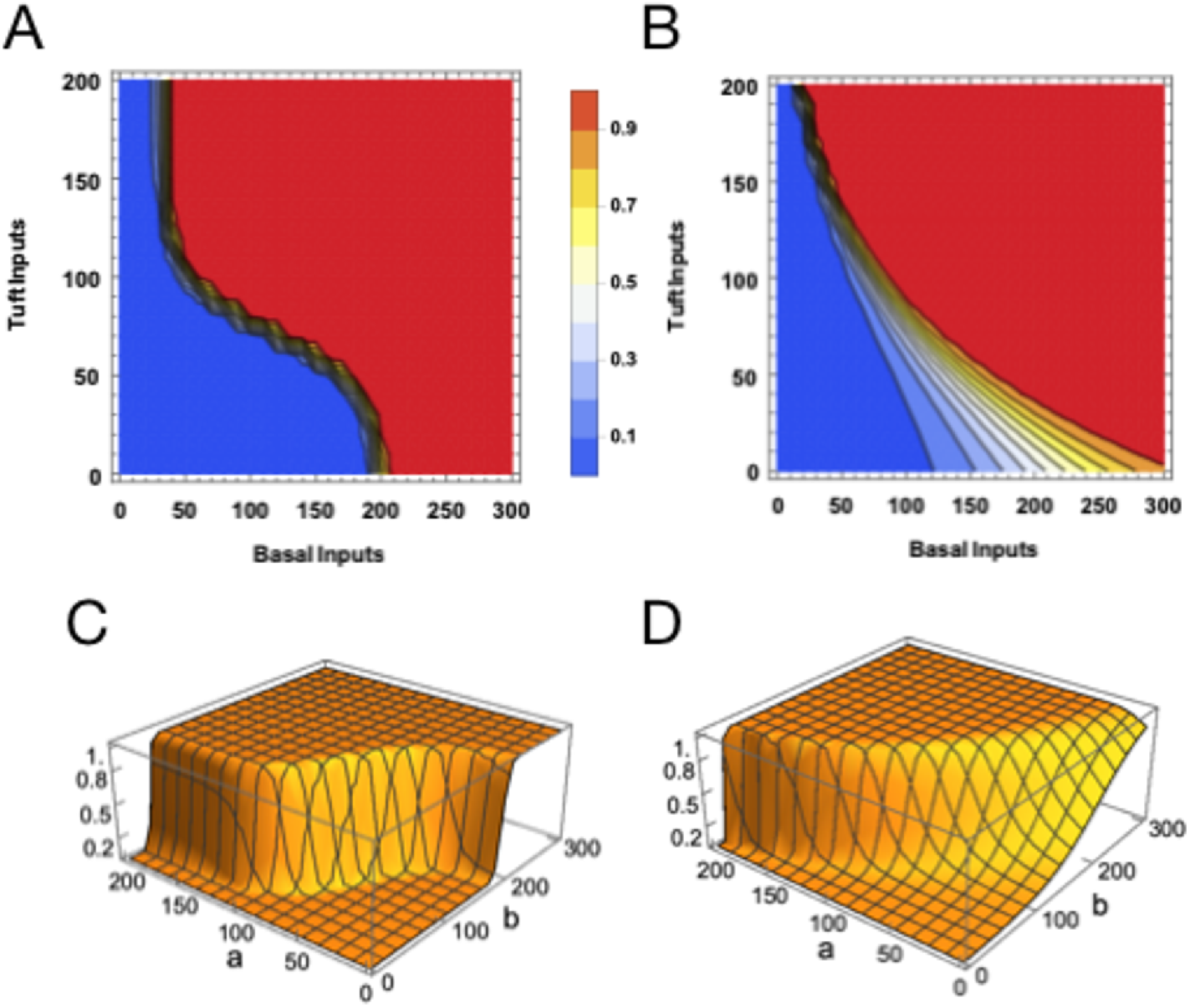
Modeling the Binary Data. **(A, B)** Contour plots for the surface of posterior predictive probability of a second AP as a function of the numbers of basal and apical tuft inputs, based on the threshold and general models, respectively. **(C, D)** Surface plots of the posterior predictive probability of a second AP as a function of the numbers of basal and apical tuft inputs, based on the threshold and general models, respectively. In each figure, the posterior predictive probabilities were computed on a 21 by 31 grid and interpolation used.

Figs 4C, 4D are surface plots of the posterior predictive probability functions. For the threshold model in Fig 4C, we notice the rather sharp transitions from almost zero probability on one side of the threshold curve to probability close to unity on the other side of the threshold curve. The general model in Fig 4D also shows such sharp transitions, especially when the apical input is large (100-200) while the basal input is low (0-100), and more gradual transitions when the apical input is lower (0-100) and the basal input is large (200-300). The two surfaces are generally similar in that for both models the sets of basal and apical inputs for which the posterior probability is close to unity, or close to zero, have a large overlap. Comparison of Fig 4C with Fig 4D shows that the posterior predictive probability surface for the general model rises more sharply for large numbers of apical and low numbers of basal input, and less sharply for lower numbers of apical input and all levels of basal input. This feature can also be noticed in the contour plots in Figs 4A, 4B.

The fit of each model to the binary response data was assessed by comparing the predictions given by the model with the actual 651 binary responses. For the threshold model, 4.2% of the responses were misclassified, whereas the error for the general model was 5.4%. Based on this posterior predictive assessment of model fit we find that the general model performs very well, and almost as well as the threshold model. The application of tenfold cross-validation in order to assess ‘out-of-sample’ prediction produced similar results: 4.3% for the threshold model and 5.5% with the general model. This similarity between ‘in-sample’ and ‘out-of-sample’ performance is due to the structure of the binary AP data.

### Application of information theory

We argue above that apical input can amplify the transmission of information about basal perisomatic input. To be made rigorous this requires quantification of information transmitted uniquely about each of the two inputs. That cannot be adequately done using classical information theory, because mutual information in that theory is defined only for a single input and a single output. There have recently been advances in the decomposition of multivariate mutual information, however, and these recent advances have now been used to specify criteria for distinguishing modulatory from driving interactions in neural systems [29]. We now apply Shannon’s classical information measures, together with partial information decompositions, to a categorised version of the action potential data produced by the detailed compartmental model reported by Shai et al. [10].

### The categorised AP data based on [10]

The output response variable, *O*, is the number of APs emitted for each combination of the basal and apical inputs. The numbers of APs range from 0 to 4, and they have been recoded into three ordinal categories, *O*_1_ – *O*_3_, containing 0-1 APs, 2 APs, 3-4 APs, respectively, since there are relatively few observations where 1 or 4 APs were obtained. The basal input was recoded into four ordinal categories: 0-60, 70-140,150-220,230-300 inputs, coded as *B*_1_ – *B*_4_, respectively. The apical input was recoded into four ordinal categories: 0-50, 60-100, 110-150, 160-200 inputs, coded as *A*_1_ – *A*_4_, respectively. This created a 4 by 4 by 3 contingency table of the recoded basal and apical inputs and the AP output. The data are displayed in Fig 5, which shows the proportions for the three AP categories for each combination of the four categories of basal input and the four categories of apical input. For the lowest category of basal input (0-60), the AP count is almost entirely 0-1 when the apical input is 0-100, but for an apical input of 110-200 we see that the proportion of observations with AP count 2-4 is about 50%. In the second lowest basal category (70-140), the AP count is 0-1 for the lowest apical category but 2-4 APs for observations in the higher apical categories (60-200). This trend from blue to red via green continues into the highest two basal categories where the AP count is 3-4 in the highest two apical categories (110-200).

**Figure 5:**
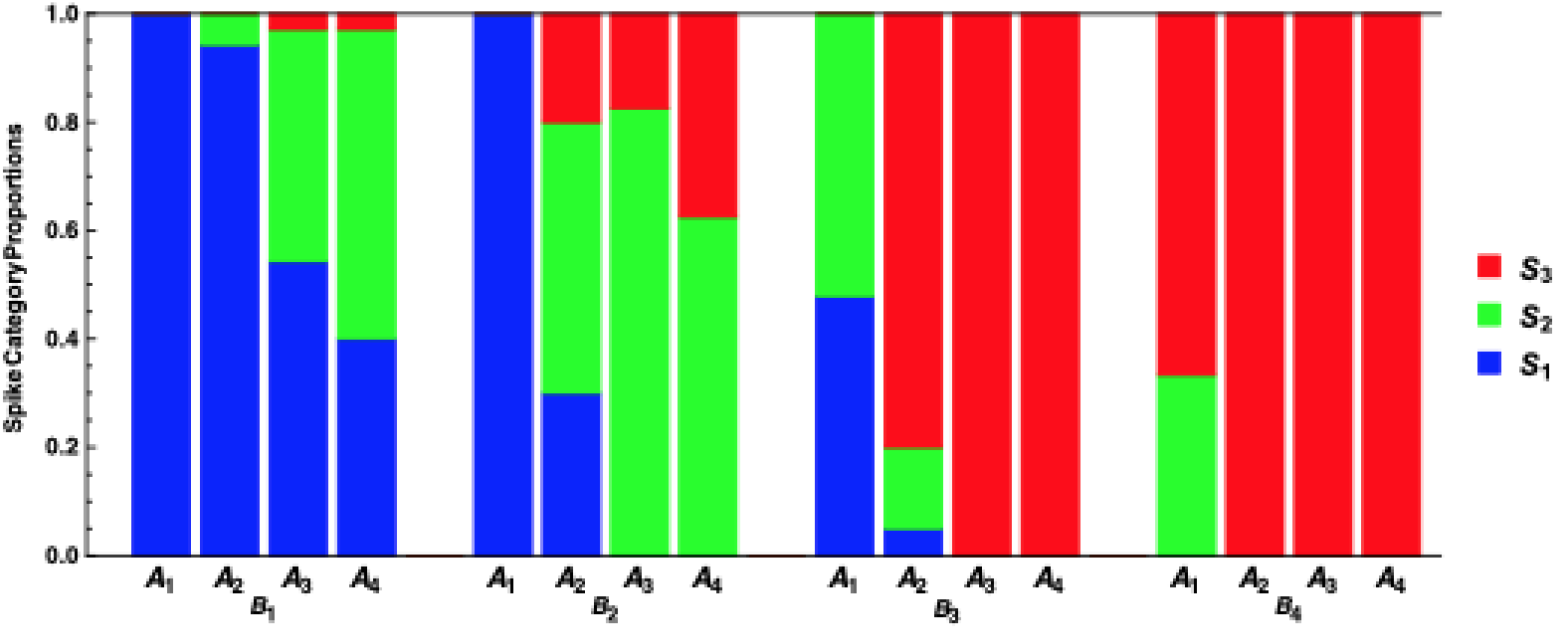
AP response distributions. A bar chart illustrating the proportion of the observations within each of the three output AP categories *O*_1_ (0-1), *O*_2_ (2), *O*_3_ (3-4) for each combination of the input basal categories *B*_1_ (0-60), *B*_2_ (70-140), *B*_3_ (150-220), *B*_4_ (230-300) and the input apical categories *A*_1_ (0-50), *A*_2_ (60-100), *A*_3_ (110-150, *A*_4_ (160-200), based on the categorised version of the AP data from [10].

### Shannon’s information measures

Using the symbol ‘*H*(*X*)’ to denote Shannon entropy of a random variable, *X*, we state some standard definitions of the measures of conditional entropy and mutual information [39,40].

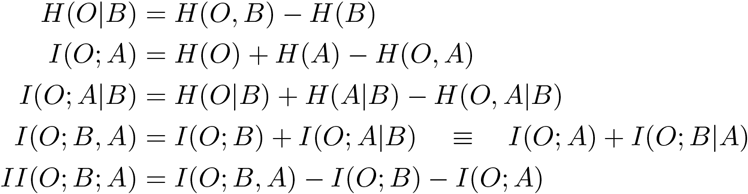

The interaction information [41], *II*(*O; B; A*), has been used as a measure of synergy [42,43].

### Application of Shannon’s information measures

The information measures were computed from these data and also the data obtained by fifteen other splittings into a 4 by 4 by 3 contingency table, and summary statistics for each measure are reported in Table 4. Details of the ‘splittings’ used are available, as referenced in the S1 File.

**Table 4:**
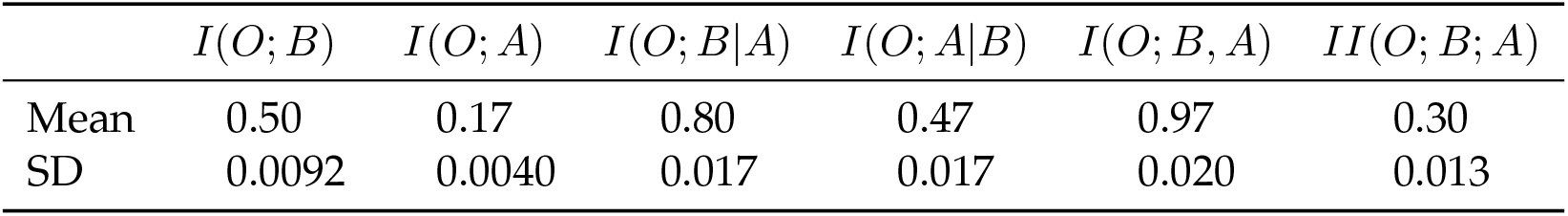
Estimated information measures. The estimated means and standard deviations (SD) of each Shannon information measure (in bits), given to two significant figures, for the sixteen categorised versions of the AP data from [10], where *O* denotes the output AP category, *B* denotes the basal input category and *A* is the apical input category. Here the output variable is the output spike category O rather than the binary output *Z*.

Based on the sample means, the joint mutual information, *I*(*O; B, A*), between the output and the bivariate distribution of the basal and apical inputs is 0.97 bits, and we notice that the mutual information, *I*(*O; B*), between the basal input and the output, 0.50 bits, is almost three times larger that the mutual information, *I*(*O; A*), between the apical input and the output, 0.17 bits. It is well known that these estimates are biased upwards [44] and so estimates of the bias were obtained. Since the number of observations (651) is large one might expect the biases to be small, and in fact they would affect only the third significant figure in each of the estimates in Table 4 if a bias correction were to be implemented.

We wish in particular to estimate the synergy in the system, since the synergistic effect of basal and tuft input is mentioned in [10]. Input from both of two distinct sources may be necessary for some transmitted information to be present. Synergy as defined within a partial information decomposition (PID) quantifies that transmission, and it plays a key role in the notion of amplification because it should be strong only when the signal being amplified is present but not strong [29].

### Application of partial information decomposition

Due to seminal work by Williams and Beer [45], it is possible to provide a finer decomposition of the information that the apical and basal inputs provide about the somatic output category. They decomposed the joint mutual information between the output category *O* and the basal and apical inputs *A, B*, considered jointly, into a sum of four non-negative terms: the shared information (Shd) that both the basal and apical inputs possess about the propagation of a somatic output category (*O*), the unique information (UnqB) that the basal input has about *O*, the unique information (UnqA) that the apical input has about *O* and the synergy (Syn) which is the information that the apical and basal inputs possess jointly about *O* that cannot be obtained by observing these two variables separately. The partial information decomposition has been applied to data in neuroscience; see, e.g. [46,47,48,49]. For a recent overview, see [50].

The basic equations [45] can be written as

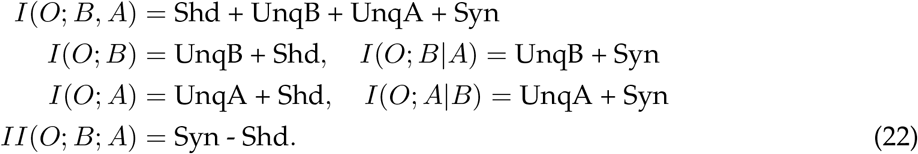

The estimate of the interaction information, *II*(*O; B; A*), reported in Table 4 is approximately 0.30 bits, and so we can deduce from Eq (22) that the estimated synergy in the system is at least 0.30 bits, on average, which is 31% of the joint mutual information *I*(*O; B, A*). The partial information decomposition [51, 52] was applied to the data and the results are given in Fig 6. The basal and apical inputs combine to transfer 45% of the joint mutual information as synergy, and they contribute to transferring 14% as shared information. There is a marked asymmetry in the estimates of the unique informations, in that the unique information due to the basal input is about ten times larger than the unique information due to the apical input. A value of zero for a unique information in an input suggests that it can contribute to the transfer of information without conveying any information about itself, and such an input is purely amplifying. Since the unique information transmitted about the apical input is close to zero (4%) this suggests that the apical input can amplify the information transmitted by the basal input in relation to the output AP category, while conveying only a very small amount of information about itself, and so is mostly amplifying and driving just a little. This lends support to the presence of apical amplification within the system; see e.g. [29]. On the other hand, the unique information transmitted about the basal input is large, at 37% of the joint mutual information, which suggests that the basal input is predominantly driving.

**Figure 6:**
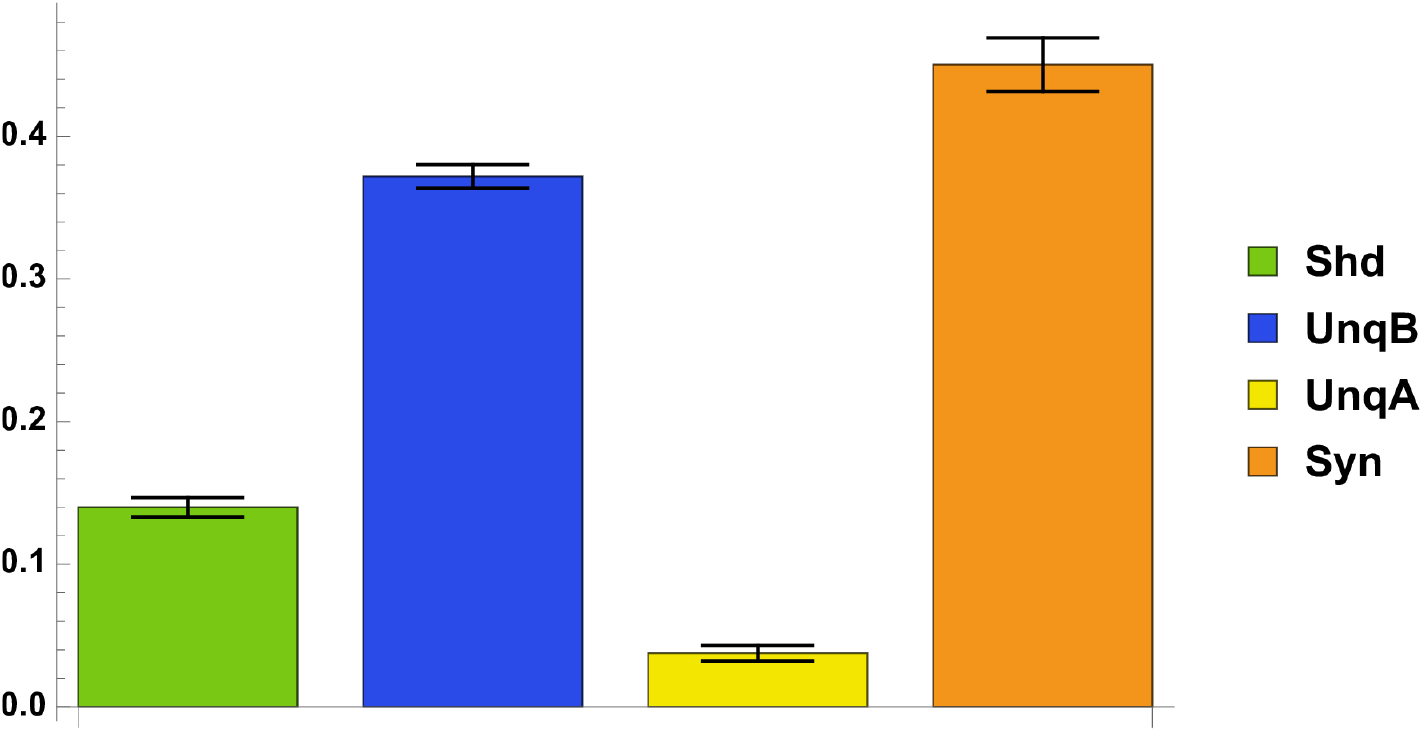
Partial information decomposition. A bar chart indicating the values of the mean shared, unique and synergistic partial information components obtained using the sixteen versions of the categorised version of the AP data from [10]. Each mean value is normalised by dividing by the mean value of *I*(*O; B, A*) and expressed as a percentage. Error bars are also shown for each component as ± one standard deviation of the sixteen values.

The results obtained with four other PIDs [45, 53, 54, 55] are provided in supplementary figure S1 Fig. The PIDs were computed using the Python package, *dit* [56].

### Alternative modes of apical function

We have so far considered the case where apical input is purely amplifying and basal input is purely driving, although intermediate cases are likely to occur. Figure 6, for example, shows that a small amount of information was transmitted uniquely about the apical input. If the apical input were purely amplifying then it would not have been small, but zero. Therefore, we now consider a wider range of cases. We first consider the unlikely, but theoretically possible, case where the functional asymmetry between apical and basal inputs is fully reversed, with apical being purely driving and basal being purely amplifying. We then consider the wide range of intermediate cases where apical and basal inputs can be partly driving and partly amplifying to various extents. Finally, we consider the case where apical and basal inputs are simply summed linearly, as is often assumed. Consideration of this wider range of cases will facilitate interpretation of any evidence of contextual feedback to basal dendrites or of feedforward input to distal apical dendrites, which may occur for various reasons to be discussed in detail elsewhere. In each of the cases considered in this section (except in Figs 7C, 7D), *P*(*S*|*a, b*) is the posterior probability, *P*(*Z* = 1|*a, b*), of an AP given apical and basal input.

**Figure 7:**
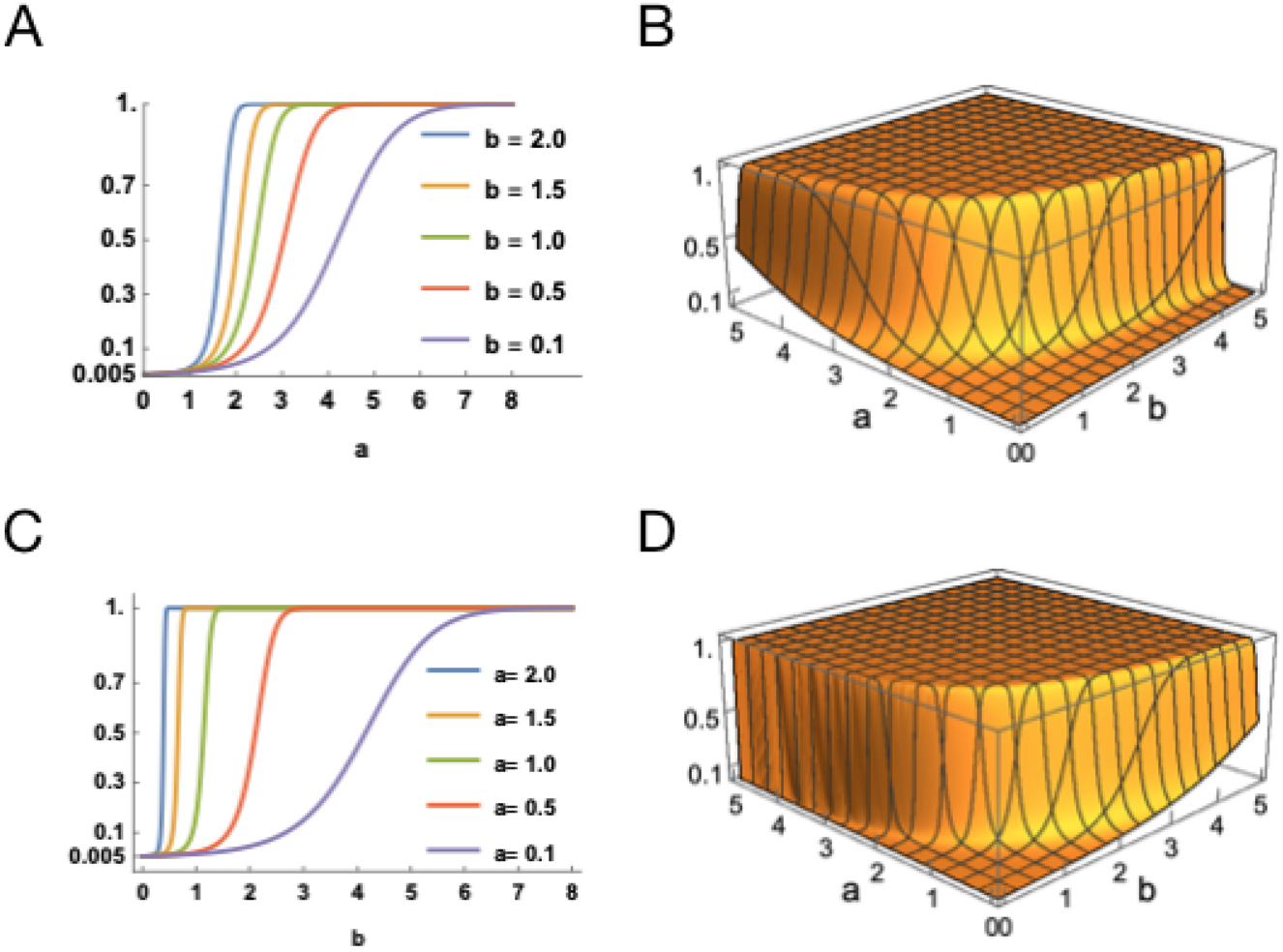
Comparing apical drive alone with basal drive. **(A)** Posterior probabilities of an AP plotted against positive values of the apical input for several values of the basal input & **(B)** a surface plot of posterior probability of an AP plotted as a function of the apical and basal inputs, both using Eqs (23), (24). **(C)** Posterior probabilities of a second AP plotted against positive values of the basal input for several values of the apical input & **(D)** a surface plot of posterior probability of a second AP plotted as a function of the apical and basal inputs, both using Eqs (12) – (14). In (C) & (D), the response probability increases rapidly when a and b are both positive, which illustrates a version of ‘coincidence detection’. The typical a priori firing probability, P(S), is assumed to be 0.005.

The first scenario is where there is basal amplification of the response to apical input when there has been no initiating bAP. Using the specific forms in Eq (10), we define the log odds and posterior probability as

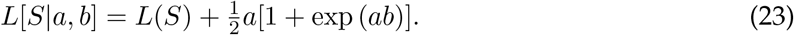

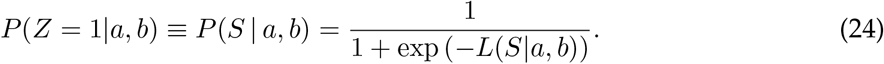

Posterior probabilities of an AP given apical drive alone are displayed in Figs 7A, 7B whereas posterior probabilities of a second AP given an initiating bAP and consequent BAC firing are shown in Figs 7C, 7D. In Fig 7A, the posterior probability of an AP is plotted as a function of positive apical input for values of the basal input ranging from 0.1 to 2.0. Even when the basal input is very weak at 0.1, we see that the probability of an action potential approaches unity when the apical input is large; the primary drive provided by the apical input is mostly responsible for this behaviour. On the other hand, when there is appreciable basal input, saturation at unity occurs very quickly for even small values of apical input. These characteristics are also evident in Fig 7C but here the probability saturates even when the basal input is less than 1. The surface plots in Figs 7B, 7D illustrate the rate at which the posterior probability saturates at unity for various values of the apical and basal inputs. The plots indicate a slight asymmetry, but for most values of a and b they are very similar, apart from the fact that the rate of saturation is more gradual in Fig 7B than in Fig 7D for lower values of a and b. Thus our weight of evidence terms suggest that basal driving coupled with BAC firing has a stronger effect than apical drive alone, especially when the level of drive is low.

The second scenario is where the amplification results from a mixture of basal and apical inputs, and we also consider the case where the basal and apical inputs combine additively. In the case of a general mixture, the forms of log odds and posterior probabilities, from (12), (23), used here are:

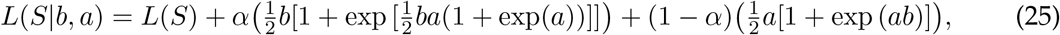

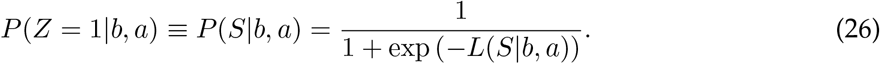

When the basal and apical inputs combine additively, the forms of log odds and posterior probabilities used are:

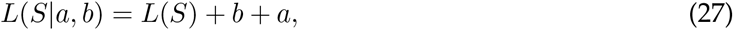

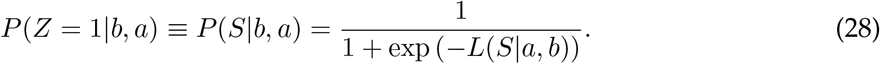

In Fig 8A, the posterior probability of an AP is plotted as a function of the basal input under an equal mixture of basal and apical amplification. The posterior probability curves saturate for small values of basal input and larger values of apical input, although larger values of basal input are required to produce saturation when the apical input is less than 1. The curves here are similar to those in Fig 7C especially when the basal input is between 0 and 1 and the apical input is between 1 and 2. These similarities given by our weight of evidence terms suggest that the probability of an AP when there is an equal mixture of basal and apical driving is similar to the probability of a second AP given an initiating bAP and consequent BAC firing, for these ranges of basal and apical input. By way of contrast, when basal and apical inputs are simply combined additively a quite different picture emerges (Fig 8C); strengthening apical input increases the probability of an action potential but has little effect on the rate at which it increases with the strength of basal input. Comparison of the surface plots in Figs 8B, 8D indicates that the presence of amplification, in contrast to linear summation, is shown by the steepening of the probability surface in Fig 8B. This distinctive effect of apical amplification has much in common with direct physiological observations reported in [57].

**Figure 8:**
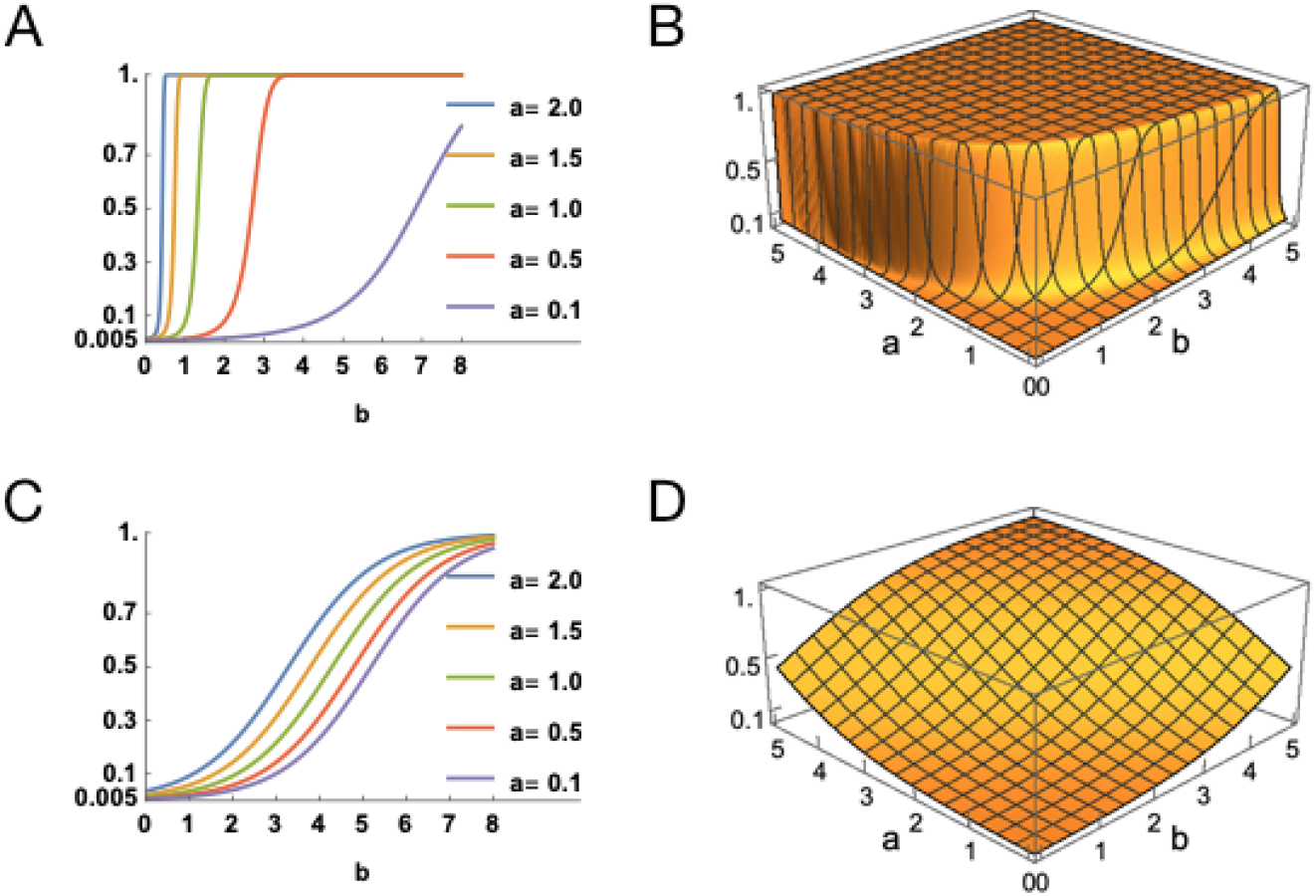
Equal mixture of basal and apical drive. **(A)** Posterior probabilities of a somatic action potential plotted against positive values of the basal input for several values of the apical input & **(B)** a surface plot of posterior probability plotted as a function of the apical and basal inputs, both using Eqs (25), (26). **(C)** Posterior probabilities of a somatic action potential, when basal and apical inputs are summed linearly, plotted against positive values of the basal input for several values of the apical input. **(D)** a surface plot of posterior probability, when basal and apical inputs are summed linearly, plotted as a function of the apical and basal inputs. Subfigures (C) and (D) are based on Eqs (27), (28). In (D), the rate of increase of the response probability is much slower than in (B). The typical baseline firing probability, *P*(*S*), is assumed to be 0.005.

### Conclusions and Discussion

The analyses presented above support the view that probabilistic inference can be context-sensitive at the level of individual neurons in particular cortical pyramidal cells. Integrate-and-fire neurons in general fire probabilistically based on knowledge stored in the strengths of their synapses. In neurons with two initiation sites and a privileged connection between them, however, the activation of one initiation site can be contingent on the other. It is this contingency that makes the two-compartmental pyramidal neurons of the cortex potentially context-sensitive and Bayesian using the principle of BAC firing. In this article, we have explored this hypothesis and conclude that BAC firing of pyramidal neurons could in principle be approximating the conditional probability that a given feature is present in the current information space given the current basal input and context. More precisely, we conclude that pyramidal neurons with BAC firing could convert the odds, *O*(feature present |basal data), into the odds, *O*(feature present | basal data and context).Useful conceptions of how probabilistic Bayesian brains combine new data with prior knowledge at the level of large neuronal circuits are already well developed [58, 59]. The importance of investing this capability in single neurons is that the cerebral cortex can potentially achieve this operation in a massively parallel fashion on all features and context simultaneously, as discussed further below.

We draw five other key conclusions from these analyses. First, BAC firing can amplify the cell’s response to basal depolarization contingent on near-coincident apical depolarization. This shows explicitly how BAC firing can be related in detail to the huge array of physiological and psychophysical evidence for contextual modulation [2, 3] in its many forms. Unlike “simple” coincidence detection, this view of apical function depends upon the marked asymmetry between the effects of apical and basal inputs that is clear in the physiological, psychophysical, and modeling data [10] and depends on the separation of the two dendritic compartments. Thus, the analyses of apical amplification presented here are consistent with previous demonstrations that contextual modulation is not a purely multiplicative interaction, which is symmetrical, but is an asymmetrical supralinear interaction [28], as shown here in Figures 2 and 6. This also implies that BAC firing approximates the activation function central to a long-standing theory of cortical computation [17,15,13] based on context-sensitive selective amplification, although there is no reason to suppose that this activation function is unique in having this property.

Second, these analyses show how amplification depends upon differentiation and communication between the two sites of integration. The somatic and apical integration sites are shown to have two functionally distinct inputs, one from outside the cell, and one from inside the cell. We have also shown that the activation functions relating the two inputs to the outputs from each site can have the same general form. This provides a clear account of the functional consequences of communication between the two sites, and provides a basis for an adequate understanding of regulation of that communication by cholinergic, adrenergic, and other modulatory systems.

Third, application of our general model to binarised AP data from a detailed multicompart-mental model [10] (Figures 3 and 4) shows that the model fits the data well, based on posterior predictive assessment, although it does not fit the data as well as the specialised threshold model. In our general model, we used particular choices of the functions, *f, g*, but it seems likely that other choices could fit the data just as well.

Fourth, application of multivariate information decomposition to categorized AP data from the detailed multicompartmental model of [10] shows that the effects of apical input approximate those expected of an amplifying interaction as defined in [29], i.e. the output transmits unique information about basal input, but little or none about apical input, even though apical input has large effects on AP output. We expect that synergy will be large only when basal input is present but weak [29, 28].

Fifth, our general decomposition can be extended to include cases where, instead of being either purely amplifying or purely driving, apical and basal inputs can each be driving and amplifying to various extents. This enables application of the analyses to neurophysiological findings providing evidence for driving effects of apical input under special conditions. This extension may play a crucial role in characterizing differences in apical function across layers, regions, species, and development. It may also help characterize changes in apical function across different states of arousal, because there is also evidence that the effects of apical input depend upon the adrenergic and cholinergic systems that regulate waking state [61], that they have a causal role in guiding overt perceptual detection [19], and that they may provide a common pathway for the effects of some general anesthetics [62, 63].

A well established method for computation in a hierarchical Bayesian network of cells is that by Lee and Mumford [27]. We haven’t used that in our Bayesian interpretation because we are concerned with the processing within a single Layer 5 pyramidal cell, whereas Lee and Mumford’s method is intended for a network of different cells. Thinking of a network of cells, the inter-cell computation could be handled using Lee and Mumford’s method while for any pyramidal cells such as those analyzed here we propose that the Bayesian interpretation described here could be used for intra-cellular processing. Thus, the two approaches are complementary.

If context-sensitive Bayesian inference is performed at the cellular level, then it becomes a privileged operation that can be carried out in a massively parallel fashion, and that will be crucial to our understanding of the information transmitted by pyramidal cells when operating in context-sensitive amplifying mode. The distinctive properties of context-sensitive two-point neurons are likely to have major implications for the short-term and long-term dynamics of the network as a whole, and though first investigated many years ago [13,14,8,64,65, ?] those implications remain largely unexplored. When this operation is hard-wired into the fabric of the cellular tissue itself it is likely to improve both speed and metabolic efficiency, and enable the cortex do things that it would not otherwise have time or energy to do. Whether context-sensitive inference at the level of the local processors also has information processing implications beyond that remains unknown.

The analyses presented here also leave many other crucial issues unresolved. As noted above, one concerns the contrast between amplifying and multiplicative interactions [28]. We assume that both forms of interaction occur in neocortex, but there are as yet few explicit empirical attempts to distinguish between them, because it is usually taken for granted that if an interaction is amplifying, then it is multiplicative [67]. That does not explain the clear functional asymmetry between somatic and apical sites of integration, however, nor does it explain how input to one site can amplify transmission of information specifically about input to the other. The contrast between amplifying and multiplicative interactions is therefore in need of thorough theoretical and empirical investigation.

Another major limitation of the current analyses is that they considered only the pyramidal neurons that, of course, exist in a complex local network – the cortical column. Although we offer a prototype mathematical description from this point of view, it is likely that the precise details of the local circuitry are ultimately vital for refining the exact description in Bayesian terms. For instance, the model could be extended to include the effects of hyperpolarizing shunting or inactivation units to the basal and apical dendrites [68, 69]. That will require clarification of the communication of inhibition between apical and somatic sites, of the time courses of recovery from inhibition, and of the local inhibitory microcircuitry. Closer attention to these issues is likely to enhance our understanding of apical function for at least three reasons. First, the types of inhibitory interneuron that target distal apical dendrites are clearly distinct from those that target basal and perisomatic regions [70, 71]. Second, there is evidence for inhibitory interneurons that amplify the output of selected pyramidal cells by specifically disinhibiting the tuft [72]. Third, there is evidence for a distinctively human class of inhibitory interneuron that targets tuft dendrites in a highly specific way [73].

One specific issue worthy of further study concerns the attractor dynamics of simple recurrent architectures in which the recurrence is provided by connections that are driving or amplifying to various extents. When analyzing the dynamics of such architectures it will be necessary to distinguish between the very fast intracellular phases of interaction between apical and somatic sites, and the slower relaxation toward an attractor that arises from intercellular interactions.

Exploration of the extent to which the idealizations studied here do or do not generalize across species, regions, development, and states of arousal has only just begun, and is likely to have far to go. The idealizations analyzed here are based on evidence that has been largely, though not wholly, collected from in vitro studies of the thick-tufted layer 5 cells of mature rodent sensory cortex that integrate activity from across all layers and provide output from the cortex at all levels of the abstraction hierarchy. Though there are grounds for supposing that they have broader relevance, these issues remain largely unexplored. Nevertheless, there are already some relevant discoveries and tantalizing hints. For example, there is evidence that somatic and apical sites are both predominantly driving in infants, with the amplifying mode of apical function becoming more available in mature animals [74, 75]. There is also evidence that apical input remains predominantly driving in the most posterior, caudal, part of mature rodent primary visual cortex (V1), whereas it is predominantly amplifying in other parts of rodent V1 [9]. Central roles for feedforward drive and feedback modulation are well established in the hierarchical regions of posterior cortex, but, as the various sub-regions of the prefrontal cortex do not seem to be orga nized into a clear hierarchy, driving and amplifying functions may be less clearly distinguished at both somatic and apical sites within prefrontal cortex. Cross-species variations in apical function may be of great importance because direct electrical recordings from the somatic and apical sites of human layer 5 pyramidal neurons show that they have enhanced electrotonic separation from the soma, as compared to those from rodents [76]. Furthermore, though there are grounds for supposing that apical function depends strongly on the state of adrenergic [61] and cholinergic arousal [77], this issue has as yet received only scant attention.

Another major unresolved issue arises from evidence that, under some circumstances, apical depolarization can drive APs in the absence of basal depolarization [57]. We therefore extended our analyses to include cases where each of the somatic and apical sites can be driving and amplifying to various extents. Intermodal effects in primary unimodal sensory regions provide clear examples of cases where apical input is clearly amplifying rather than driving. For example, anatomical and physiological evidence indicates that the effects of auditory information on pyramidal cells in V1 sharpens their selective sensitivity and raises the salience of their output via the apical dendrites in layer 1 [78]. Nevertheless, as noted above, the balance between driving and amplifying effects of apical input may vary even within a single neocortical region such as V1 [9]. There is some hint of a sharp transition from a driving to an amplifying effect of apical depolarization as apical trunk-length increases in V1 [9], but whether there is a clear dichotomy or a continuum between these two forms of apical function in other regions remains unknown. Much remains to be discovered concerning the conditions under which apical input is purely amplifying, purely driving, or a mixture of the two. Whatever the resolution to those issues, however, if some pyramidal cells do indeed often function as two-point processors, then system-level network diagrams showing inputs to pyramidal neurons represented as single undifferentiated points of integration, though abundant, are grossly underspecified.

Grounding context-sensitive selective amplification in subcellular processes may have far-reaching implications not only for cognition [79, 62, 14, 64, 65] but also for psychopathology. It may have relevance to psychopathology because there is evidence that apical malfunction is involved in pathologies as diverse as schizophrenia [80, 81], autoimmune anti-NMDAR encephalitis [80, 82], absence epilepsy [83], and foetal alcohol spectrum disorder [84, 85]. If so, then studies of apical function may cast light on those disorders, and those disorders may cast light on apical function. For in-depth discussion of impaired context-sensitivity in schizophrenia see [86]. For detailed compartmental modelling of ion channels by which risk genes for schizophrenia could impair apical function see [87].

The evidence that some neocortical neurons can function as two-point processors raises the possibility of using two-point processors to approximate some powerful machine learning algorithms while making them more biologically plausible. This has already been explored to some extent [88, 89, 90], but the use of apical input to amplify current response to basal depolarization may not only make such algorithms more biologically plausible, but may also enhance their capabilities. As contextual amplification does not compromise the flow of uniquely feedforward information, amplifying input could come from diverse sources, as it does in neocortex. Consider the exceptionally well-known hierarchical feedforward architecture of [91], for example. At the convolutional levels of that architecture contextual input to each local processing element could include information from other streams at the same hierarchical level. It could also include information about goal states specified for higher levels in the hierarchy, and that may be mediated via intermediate levels, or be specified by direct feedback connections that skip levels, analogous to those known to exist in neocortex [92]. That would remove one major difference between deep learning algorithms and neocortex because learning in neocortex is usually the consequence of effects on current processing, whereas back-propagation in deep learning algorithms effects learning but not processing of the current data. For initial studies of learning and processing in nets with many parallel streams of processing composed of two-point local processors with context sensitivity see [13], and in nets with feedback from a higher level see [66]. For approximations by which the learning rules used in those studies can be scaled-up see [15]. For discussion of how that form of learning and processing may advance beyond the pioneering work of [93] see [94]. Though the development of any such enhanced form of machine learning algorithm would not prove that the amplifying mode of apical function makes a crucial contribution to mental life, it would certainly show that it could do so, and as a consequence greatly increase neurobiological interest in these possibilities.

In summary, we conclude that neocortical pyramidal neurons with two functionally distinct points of integration can plausibly be described as operating in a context-sensitive Bayesian way on two distinct information streams. The key aspect of their biophysics is the privileged connection (the apical dendrite) that connects these two separate sites of integration. In this way, the operation of each of these sites can be contingent on the other in a way that enables context-sensitive Bayesian inferences that would otherwise be far more complex to realize at the network level.

## Methods

### A Bayesian interpretation of intra-site computation

Following the applications of Bayes’ theorem in the Results section, we provide derivations of odds and log odds and discussion of the weight of evidence terms. The prior odds and log odds in favour of a second AP are

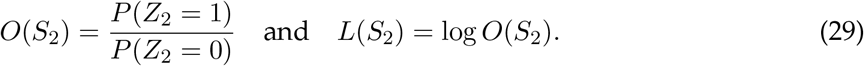

Using (1), the posterior odds and log odds in favour of a second AP given the basal input are

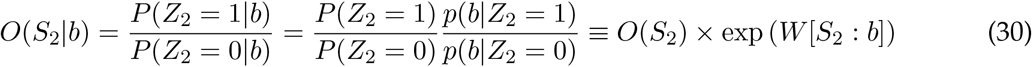

and

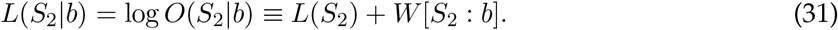

Here the weight of evidence term is

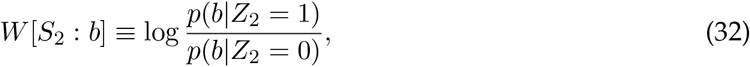

which is the logarithm of the Bayes factor in favour of event *S*_2_ provided by the basal input, *b*. This Bayes factor is a ratio of the likelihood of a second AP, given the basal input, and the likelihood of no second AP, given the basal input. A value greater than 1 for the Bayes factor would favor the event *S*_2_, as opposed to the complementary event 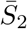, and this corresponds to a positive value for the weight of evidence term.

Similarly, we can use Eq (4) to update the posterior odds in Eq (30) once the apical input is observed in addition to the basal input:

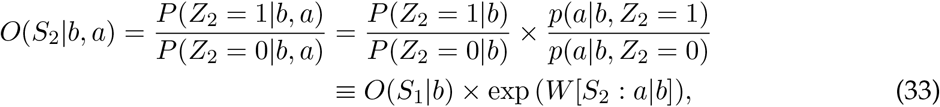

where the weight of evidence term is

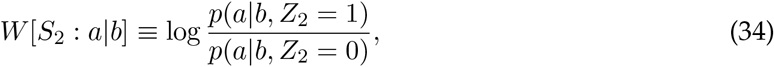

which is the Bayes factor in favor of a second AP provided by the the apical input, *a*, given the basal input, *b*. This Bayes factor is a ratio of the likelihood of a second AP, given both the basal and apical inputs, and the likelihood of no second AP, given both inputs. A value greater than 1 for this Bayes factor would favor the occurrence of a second AP, as opposed to its non-occurrence, and this corresponds to a positive value for the weight of evidence term. Applying logarithms in Eq (33), we have

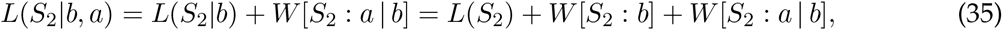

using Eq (31).

### Activation functions used in the apical and somatic sites

The functional forms in (10) are used in both sites of integration. In the AIS, the apical input is driving and the occurrence of a bAP is amplifying (or attenuating). Therefore, we take *x* = *a* and *y* = *z*_1_, so that the apical activation in Phase 1 is just *f* (*a*) = *a*, and the apical activation during Phase 2 is 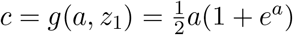.

In the SIS, the basal input is driving and the apical activation is amplifying (or attenuating). Therefore, in the SIS where we are principally interested in Phase 2, we take *x* = *b* and *y* = *c* ≡ *f* (*a*) + *g*(*a, z*_1_) in (10), which results in the somatic activation during Phase 2 becoming

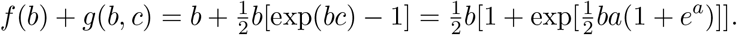

The functions *f, g* in (10) are similar to those used in previous work on artificial neural nets [66, 13] and more recently in work on partial information decomposition [29, 28]. Further details regarding these functions are given in the supplementary file S3 File.

### Monte Carlo approximation of posterior predictive probabilities

We describe the formulae for the general model; they are analogous for the threshold model.

The posterior predictive probabilities in Eqs (18), (21) were computed using the output from `rstan` as Monte Carlo approximations, using (36). The posterior predictive probability in Eq (21) can be written as a posterior expectation, as follows:

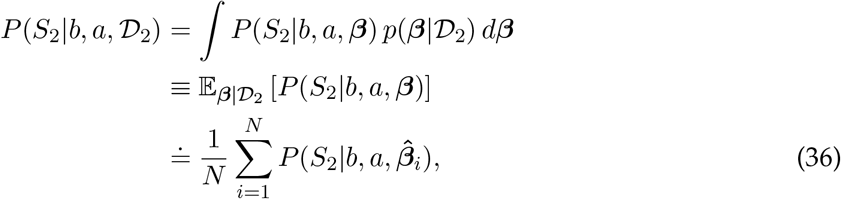

where 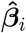 is the estimate of *β* generated in the ith of the *N* simulations. For further detail, see the files referenced in the S1 File.

## Supporting information

**S1 File. Data analysis**. Details of the modeling and analysis, together with R, RStan, Python and Mathematica code.

**S2 File Bayesian interpretation** An illustration of the implementation.

**S3 File Choices of activation functions** *f, g*. Details of the derivation and an application.

**S1 Fig Results from other PIDs**. Bar charts indicating the values of the mean shared, unique and synergistic partial information components obtained using the sixteen versions of the categorised version of the AP data from [10]. Each mean value is normalised by dividing by the mean value of *I*(*O; B, A*) and expressed as a percentage. Error bars are also shown for each component as ± one standard deviation of the sixteen values. **A** Imin [45], **B** Iproj[53], **C** Iccs [54], **D** Idep [55]

## Acknowledgments

We are grateful to the editors and three anonymous reviewers for helpful comments on an earlier version of the manuscript which led to improvements in both the organisation and the content.

