## Supplementary File S1 for "Bayesian modeling of BAC firing as a mechanism for apical amplification in neocortical pyramidal neurons"

### Data Analysis

#### Action Potential Data from Shai et al. [10]

The data from Shai et al. (2015) that is used in the Results section is available in the file spikes_.dat at:

https://senselab.med.yale.edu/ModelDB/ShowModel.cshtml?model=180373&
file=/ShaiEtAl2015/data/spikes_.dat#tabs-2

#### GitHub repository

The files described in the sequel are available from the GitHub repository:

https://github.com/JWKay/Pyramid.

#### Binarised action potential data: determining the thresholds

The action potential (AP) data were binarised, as described in the R script Figures1.R, and saved in the file spbin.csv. For each of the 21 numbers of tuft inputs, a penalized binary logistic regression model was applied, making use of the R package *logistf*. The data exhibit *complete separation*, so a form of regularization is required; otherwise the maximum likelihood estimates would grow without bound.

For each value of the number of tuft inputs, the explanatory variable, $b_i$, was the number of basal inputs ($i = 1, 2, \ldots, 31$), and the binary response, $z_i$, was 1 if a second AP was recorded, and 0 otherwise. A penalized binary logistic regression was performed, with $Z_i$ following a Bernoulli distribution with probability $p_i$ of a second AP when the number of basal inputs is $b_i$, and linear predictor,

$$\log \frac{p_i}{1 - p_i} = \alpha_1 + \alpha_2 b_i,$$

where $\alpha_1, \alpha_2$ are unknown parameters. The estimated threshold was computed as $-\alpha_1/\alpha_2$, the value of basal input for which the probability of a second AP is 0.5, and the estimated standard error computed using the delta method. This produced an estimate of the threshold for each of the 21 values of the number of tuft inputs. The corresponding pairs of number of tuft inputs and the threshold were saved in the file threshdat.csv, for use in Figure 3A. The Stan model code used for making posterior predictive predictions of the thresholds is given in BayNRpred.stan. The predicted thresholds, together with pointwise 95% prediction limits, were stored in the file pred.csv for use in producing Figure 3B.

#### Threshold Model for the binarised action potential data

As described in the Bayesian Modeling subsection of the Methods section, a weighted Bayesian nonlinear regression was fitted to the threshold data, by making use of the *rstan* package. The details are provided in the R script Figures1.R. The Stan model is given in BayNR.stan. A seed and initial values are included in the calling code to ensure that the results are reproducible. Four Markov chains were run for 3000 iterations, with the first 500 iterations in each chain being discarded as 'warmup'. The usual convergence checks were conducted. 10,000 samples were stored for use in computing Monte Carlo approximations of the posterior predictive probabilities. Their values, along with numbers of basal and apical inputs, were saved in the file TMdat.csv, which was used in Mathematica to produce Figs 4A, 4C, 4E. Details of the posterior predictive assessments, including the code for tenfold crossvalidation, are also given in the file Figures1.R

#### General Model for the binarised action potential data

The details of the general model are provided in the Bayesian Modeling subsection of the Methods section, with R script Figures2.R containing the computational details. The Stan model is defined in the file GenMod.stan. A seed and initial values are included in the calling code to ensure that the results are reproducible. Four chains were run for 3000 iterations, with the first 500 iterations in each chain being discarded as 'warmup'. 10,000 samples were stored for use in computing Monte Carlo approximations of the posterior predictive probabilities. Their values, along with numbers of basal and apical inputs, were saved in the file GMdat.csv, which was used in Mathematica to produce Figs 4B, 4D. Details of the posterior predictive assessments, including the code for tenfold crossvalidation, are also given in the file Figures2.R.

#### Analysis of the categorized action potential data

The action potential data from Shai et al. [10] was categorized, as described in the Results section and as shown in the R script Figures3.R. The categorized data were then used in the Python 3.5 script cspid.py in order to compute estimates of classical Shannon measures along with five different partial information decompositions, yielding the results given in Figure 5. In further work, a further fifteen splits of the binary data were also produced, as described in the R script Figures4.R. These data were then processed using the Python script cspid2.py and the results written to the file PIDout.txt, which was read into R for computation of the results which appear in Table 4 and Figure 6.

#### Illustrative example, Supplementary file S2

An illustrative example of how to implement the Bayesian interpretation of BAC firing is provided in the supplementary file, S2 File. The computation was produced using the R script Supp-S2-BN.

#### Plots of Figures 2-8, Supplementary Material

Plots of figures 2-8, two figures in the supplementary file, S2 File, and the supplementary figure S1 Fig were produced using Mathematica, and the full code and plots are provided in the computable document format file KPAGLplots.cdf. This can be viewed in a cdf player that can be obtained free of charge for devices (as an app) and desktops from Wolfram at

https://www.wolfram.com/cdf-player/

The details are also available in portable document format as KPAGLplots.pdf.
