## Supplementary File S2 for "Bayesian modeling of BAC firing as a mechanism for apical amplification in neocortical pyramidal neurons"

### An illustration of the Bayesian interpretation of BAC firing

#### Introduction

In the absence of real data we will use simulated data in this illustration. By assuming that the numbers of basal and apical inputs are large and not all too strongly correlated, we will assume Gaussian probability models for the various generative models for the integrated basal and apical inputs, respectively, by a version of the Central Limit Theorem. Standard Bayesian distributional results are used to define a noninformative prior for the unknown parameters and also to find their posterior distribution given the data. We then define the probability of a second action potential given the unknown parameters. This will be estimated by using Monte Carlo simulation from the posterior distribution. In addition, the Monte Carlo standard error will be estimated. We also will compute 90% equal-tailed credible intervals for the unknown probability of a second action potential. The various standard Bayesian results quoted in the sequel are taken from the book, Bayesian Data Analysis, 3rd edition, by Gelman et al. (Section 3.2, Section 14.2, p. 583).

#### The Data

Data are available in the form $\{z_i, b_i, a_i : \ i = 1, \ldots, n\}$. Since we will be dealing with generative models of the basal input given that $Z_2 = 0, 1$, respectively, as well as for the apical input given the basal input and $Z_2 = 0, 1$, respectively, we can consider the data in the form $\mathcal{D} = \{\mathbf{a}_0, \mathbf{b}_0, \mathbf{a}_1, \mathbf{b}_1\}$, where the $\mathbf{a}_i, \mathbf{b}_i$ are the apical and basal inputs, respectively when $Z_2 = i, (i = 0, 1)$. The random variable $Z_2$ records whether or not a second action potential (AP) occurs. We let $n_0$ be the number of observations for which $Z_2 = 0$ and $n_1$ be the number of observations for which $Z_2 = 1$. So, $n = n_0 + n_1$.

#### Generative models for basal and apical inputs

We let the random variables, $A_i, B_i$ describe the apical and basal input, respectively, when $Z_2 = i$. First we define generative models for $B_0, B_1$ given that $Z_2 = 0, 1$, respectively. We assume that $B_0$ is Gaussian with mean $\mu_0$ and variance $\sigma_0^2$, and that $B_1$ is Gaussian with mean $\mu_1$ and variance $\sigma_1^2$. Therefore, probability density functions (pdf) of $B_0, B_1$ are:

$$p(b_0|\mu_0, \sigma_0^2) = \frac{1}{\sqrt{2\pi}\sigma_0} \exp\left(-\frac{1}{2\sigma_0^2}(b_0 - \mu_0)^2\right), \tag{1}$$

$$p(b_1|\mu_1, \sigma_1^2) = \frac{1}{\sqrt{2\pi}\sigma_1} \exp\left(-\frac{1}{2\sigma_1^2}(b_1 - \mu_1)^2\right). \tag{2}$$

The generative model for $A_0$ given that $B_0 = b_0$ and $Z_2 = 0$ is a Gaussian linear regression model with mean $\alpha_0 + \beta_0 b_0$ and variance $\sigma_3^2$. Similarly, the generative model for $A_1$ given that $B_1 = b_1$ and $Z_2 = 1$ is a Gaussian linear regression model with mean $\alpha_1 + \beta_1 b_1$ and variance $\sigma_4^2$. The conditional pdfs for $A_i$ given $B_i = b_i$ and $Z_2 = i$ are as follows.

$$p(a_0|b_0, \alpha_0, \beta_0, \sigma_3^2) = \frac{1}{\sqrt{2\pi}\sigma_3} \exp\left(-\frac{1}{2\sigma_3^2}(a_0 - \alpha_0 - \beta_0 b_0)^2\right), \tag{3}$$

$$p(a_1|b_1, \alpha_1, \beta_1, \sigma_4^2) = \frac{1}{\sqrt{2\pi}\sigma_4} \exp\left(-\frac{1}{2\sigma_4^2}(a_1 - \alpha_1 - \beta_1 b_1)^2\right). \tag{4}$$

Let the vector $\boldsymbol{\theta}$ contain all the unknown parameters, so that

$$\boldsymbol{\theta} = \{\mu_0, \sigma_0^2, \mu_1, \sigma_1^2, \alpha_0, \beta_0, \sigma_3^2, \alpha_1, \beta_1, \sigma_4^2\}$$

#### Prior to Posterior

We assume *a priori* that the unknown parameters in $\boldsymbol{\theta}$ are mutually independent, and we assign noninformative priors for each of the four sets of parameters (Gelman, p. 64, p. 355) as follows.

$$p(\mu_0, \sigma_0^2) \propto (\sigma_0^2)^{-1}, \quad p(\mu_1, \sigma_1^2) \propto (\sigma_1^2)^{-1}$$
$$p(\alpha_0, \beta_0, \sigma_3^2) \propto (\sigma_3^2)^{-1}, \quad p(\alpha_1, \beta_1, \sigma_4^2) \propto (\sigma_4^2)^{-1}$$

The likelihood function of the parameter $\boldsymbol{\theta}$ given the data $\mathcal{D}$ is

$$p(\mathcal{D}|\boldsymbol{\theta}) = \prod_{i=1}^{n_0} p(b_{0i}|\mu_0, \sigma_0^2) \prod_{i=1}^{n_0} p(a_{0i}|b_{0i}, \alpha_0, \beta_0, \sigma_3^2) \prod_{i=1}^{n_1} p(b_{1i}|\mu_1, \sigma_1^2) \prod_{i=1}^{n_1} p(a_{1i}|b_{1i}, \alpha_1, \beta_1, \sigma_4^2) \tag{5}$$

in factorized form. Since the likelihood factorizes and the prior factorizes we find that the posterior distribution of $\boldsymbol{\theta}$ given the data $\mathcal{D}$ also factorizes:

$$p(\boldsymbol{\theta}|\mathcal{D}) = p(\mu_0, \sigma_0^2|\mathbf{b}_0) p(\alpha_0, \beta_0, \sigma_3^2|\mathbf{a}_0, \mathbf{b}_0) p(\mu_1, \sigma_1^2|\mathbf{b}_1) p(\alpha_1, \beta_1, \sigma_4^2|\mathbf{a}_1, \mathbf{b}_1) \tag{6}$$

which means that the four sets of parameters in the posterior distributions are mutually independent given the data $\mathcal{D}$. Before stating the forms of the respective posterior distributions, we define the summary statistics of the data that are required. First we provide expression for sample means and variances.

$$\bar{b}_0 = \frac{1}{n_0} \sum_{i=1}^{n_0} b_{0i} \tag{7}$$

$$\bar{b}_1 = \frac{1}{n_1} \sum_{i=1}^{n_1} b_{1i} \tag{8}$$

$$s_0^2 = \frac{1}{n_0 - 1} \sum_{i=1}^{n_0} (b_{0i} - \bar{b}_0)^2 \tag{9}$$

$$s_1^2 = \frac{1}{n_1 - 1} \sum_{i=1}^{n_1} (b_{1i} - \bar{b}_1)^2 \tag{10}$$

Here, $b_{0i}$ is the $i$th observation of the basal input when $Z_2 = 0$, and $b_{1i}$ is the $i$th observation of the basal input when $Z_2 = 1$, with $\bar{b}_0, \bar{b}_1$ denoting the sample means. Also required are summary statistics from the simple linear regressions. We let $\psi_0 = \{\alpha_0, \beta_0\}$ and $\psi_1 = \{\alpha_1, \beta_1\}$. Then

$$\hat{\psi}_0 = (X_0^T X_0)^{-1} X_0^T \mathbf{a}_0, \tag{11}$$

$$\hat{\psi}_1 = (X_1^T X_1)^{-1} X_1^T \mathbf{a}_1, \tag{12}$$

$$V_0 = (X_0^T X_0)^{-1}, \tag{13}$$

$$V_1 = (X_1^T X_1)^{-1}, \tag{14}$$

and

$$X_0 = \begin{bmatrix} 1 & b_{01} \\ 1 & b_{02} \\ \vdots & \vdots \\ 1 & b_{0n_0} \end{bmatrix}, \quad X_1 = \begin{bmatrix} 1 & b_{11} \\ 1 & b_{12} \\ \vdots & \vdots \\ 1 & b_{1n_1} \end{bmatrix}.$$

Also

$$s_3^2 = \frac{1}{n_0 - 2} (\mathbf{a}_0 - X_0 \hat{\psi}_0)^T (\mathbf{a}_0 - X_0 \hat{\psi}_0), \quad s_4^2 = \frac{1}{n_1 - 2} (\mathbf{a}_1 - X_1 \hat{\psi}_1)^T (\mathbf{a}_1 - X_1 \hat{\psi}_1).$$

The standard forms of the posterior distributions are now stated.

$$\sigma_0^2 | \mathbf{b}_0 \sim \text{Inv-}\chi^2(n_0 - 1, s_0^2), \quad \mu_0 | \sigma_0^2, \mathbf{b}_0 \sim N(\bar{b}_0, \sigma_0^2 / n_0) \tag{15}$$

$$\sigma_1^2 | \mathbf{b}_1 \sim \text{Inv-}\chi^2(n_1 - 1, s_1^2), \quad \mu_1 | \sigma_1^2, \mathbf{b}_1 \sim N(\bar{b}_1, \sigma_1^2 / n_1) \tag{16}$$

$$\sigma_3^2 | \mathbf{a}_0, \mathbf{b}_0 \sim \text{Inv-}\chi^2(n_0 - 2, s_3^2), \quad \psi_0 | \mathbf{a}_0, \mathbf{b}_0 \sim BN(\hat{\psi}_0, V_0 \sigma_3^2) \tag{17}$$

$$\sigma_4^2 | \mathbf{a}_1, \mathbf{b}_1 \sim \text{Inv-}\chi^2(n_1 - 2, s_4^2), \quad \psi_1 | \mathbf{a}_1, \mathbf{b}_1 \sim BN(\hat{\psi}_1, V_1 \sigma_4^2) \tag{18}$$

Here, Inv-$\chi^2$ denotes the inverse chi-squared distribution, while $N, BN$ denote the univariate Gaussian and bivariate Gaussian distributions, respectively. Noinformative priors have been assumed but here, as long as $n_0 > 2$ and $n_1 > 2$, and $X_0, X_1$ both have full rank 2, then the posterior distributions will be perfectly proper. This is the case in this illustration.

#### Simulation from the posterior distribution

The posterior distributions in Eqs (15-18) are all standard and so the Monte Carlo method can be used to generate random draws from the posterior distribution, given the values of the statistics in Eqs (7-14), and there is no need to resort to MCMC methods.

First we generate the pair $(\sigma_0^2, \mu_0)$.
Draw $X_0$ from a $\chi^2(n_0 - 1)$ distribution, and take $\sigma_0^2 = (n_0 - 1)s_0^2 / X_0$.
Using this generated value of $\sigma_0^2$, then draw $\mu_0$ from $N(\bar{b}_0, \sigma_0^2 / n_0)$.

Then we generate the pair $(\sigma_1^2, \mu_1)$.
Draw $X_1$ from a $\chi^2(n_1 - 1)$ distribution, and take $\sigma_1^2 = (n_1 - 1)s_1^2 / X_1$.
Using this generated value of $\sigma_1^2$, then draw $\mu_1$ from $N(\bar{b}_1, \sigma_1^2 / n_1)$.

Then generate $(\sigma_3^2, \psi_0)$.

Draw $X_3$ from a $\chi^2(n_0-2)$ distribution, and take $\sigma_3^2 = (n_0-2)s_3^2/X_3$.
Then use $\sigma_3^2$ in drawing $\psi_0 = \{\alpha_0, \beta_0\}$ from the bivariate Gaussian distribution with mean vector $\hat{\psi}_0$ and covariance matrix $V_0\sigma_3^2$.

Then generate $(\sigma_4^2, \psi_1)$.
Draw $X_4$ from a $\chi^2(n_1-2)$ distribution, and take $\sigma_4^2 = (n_1-2)s_4^2/X_4$.
Then use $\sigma_4^2$ in drawing $\psi_1 = \{\alpha_1, \beta_1\}$ from the bivariate Gaussian distribution with mean vector $\hat{\psi}_1$ and covariance matrix $V_1\sigma_4^2$.

#### Posterior predictive probability and credible interval

Let $a, b$ be new (or future) values of apical, basal input, respectively. Then we wish to estimate the probabilities of a second AP (the event $S_2$)

$$P(S_2|b,\boldsymbol{\theta}), \quad P(S_2|b,a,\boldsymbol{\theta}),$$

given the basal input $b$ and both basal and apical input $b, a$, respectively. These probabilities are functions of the unknown parameters, $\boldsymbol{\theta}$, although the first probability is a function of only the first four components of $\boldsymbol{\theta}$. For simplicity, we employ the same vector $\boldsymbol{\theta}$ in both cases. Now

$$P(S_2|b,\boldsymbol{\theta}) = \frac{1}{1+\exp\left(-L(S_2|b,\boldsymbol{\theta})\right)},$$

$$P(S_2|b,a,\boldsymbol{\theta}) = \frac{1}{1+\exp\left(-L(S_2|b,a,\boldsymbol{\theta})\right)},$$

where $L(S_2|b,\boldsymbol{\theta}), L(S_2|b,a,\boldsymbol{\theta})$ are the log odds ratios given by

$$\begin{aligned} L(S_2|b,\boldsymbol{\theta}) &= L(S_2) + W[S_2 : b|\boldsymbol{\theta}] \\ L(S_2|b,a,\boldsymbol{\theta}) &= L(S_2) + W[S_2 : b|\boldsymbol{\theta}] + W[S_2 : a|b,\boldsymbol{\theta}] \end{aligned}$$

respectively. These expressions involves weight of evidence (WoE) terms, now expressed also in terms of the unknown parameters $\boldsymbol{\theta}$, and $L(S_2)$ is the prior log odds. By Eqs (1-4), the WoE terms are

$$W[S_2 : b|\boldsymbol{\theta}] = \log\frac{p(b_1|\mu_1,\sigma_1^2)}{p(b_0|\mu_0,\sigma_0^2)} = \log\frac{\sigma_0}{\sigma_1} - \frac{1}{2\sigma_1^2}(b-\mu_1)^2 + \frac{1}{2\sigma_0^2}(b-\mu_0)^2$$

and

$$W[S_2 : a|b,\boldsymbol{\theta}] = \log\frac{p(a_1|b_1,\alpha_1,\beta_1,\sigma_4^2)}{p(a_0|b_0,\alpha_0,\beta_1,\sigma_3^2)} = \log\frac{\sigma_3}{\sigma_4} - \frac{1}{2\sigma_4^2}(a-\alpha_1-\beta_1 b)^2 + \frac{1}{2\sigma_3^2}(a-\alpha_0-\beta_0 b)^2$$

The posterior predictive probability of event $S_2$ given a value for $b$ and the data is equal to

$$P(S_2|b,\mathcal{D}) = \int P(S_2|b,\boldsymbol{\theta})p(\boldsymbol{\theta}|\mathcal{D})\,d\boldsymbol{\theta}. \tag{19}$$

The posterior predictive probability in Eq (19) can be written as a posterior expectation, as follows:

$$
\begin{aligned}
P(S_2|b,\mathcal{D}) &= \int P(S_2|b,\boldsymbol{\theta})\, p(\boldsymbol{\theta}|\mathcal{D})\, d\boldsymbol{\theta} \\
&\equiv \mathbb{E}_{\boldsymbol{\theta}|\mathcal{D}}\left[P(S_2|b,a,\boldsymbol{\theta})\right] \\
&\doteq \frac{1}{N}\sum_{i=1}^{N} P(S_2|b,a,\hat{\boldsymbol{\theta}}_i),
\end{aligned}
$$

where $\hat{\boldsymbol{\theta}}_i$ is the estimate of $\boldsymbol{\theta}$ generated in the $i$th of the $N$ simulations. Thus, it can be estimated by the Monte Carlo method.

Similarly, the posterior predictive probability of event $S_2$ given values for $a, b$ and the data is equal to

$$
P(S_2|b,a,\mathcal{D}) = \int P(S_2|b,a,\boldsymbol{\theta})p(\boldsymbol{\theta}|\mathcal{D})\, d\boldsymbol{\theta}. \tag{20}
$$

The posterior predictive probability in Eq (20) can be written as a posterior expectation, as follows:

$$
\begin{aligned}
P(S_2|b,a,\mathcal{D}) &= \int P(S_2|b,a,\boldsymbol{\theta})\, p(\boldsymbol{\theta}|\mathcal{D})\, d\boldsymbol{\theta} \\
&\equiv \mathbb{E}_{\boldsymbol{\theta}|\mathcal{D}}\left[P(S_2|b,a,\boldsymbol{\theta})\right] \\
&\doteq \frac{1}{N}\sum_{i=1}^{N} P(S_2|b,a,\hat{\boldsymbol{\theta}}_i),
\end{aligned}
$$

where $\hat{\boldsymbol{\theta}}_i$ is the estimate of $\boldsymbol{\theta}$ generated in the $i$th of the $N$ simulations. Thus, it also can be estimated by the Monte Carlo method.

These expression give point estimates of the posterior probabilities in (), (), but the probabilities are parametric functions of $\boldsymbol{\theta}$ and so given the posterior distribution of $\boldsymbol{\theta}$ it is possible to estimate other quantities of the posterior distributions of these probabilities by using the parameters simulated from the posterior distribution, $p(\boldsymbol{\theta}|\mathcal{D})$. We do this only for $P(S_2|b,a,\boldsymbol{\theta})$.

In particular we can estimate the Monte Carlo standard error of the estimate in (20) as the standard deviation of $P(S_2|b,a,\boldsymbol{\theta})$ divided by $\sqrt{N}$. We can also find a 90% equal-tailed credible interval for $P(S_2|b,a,\boldsymbol{\theta})$ by finding the 0.05th and 0.95th quantiles of the posterior distribution of $P(S_2|b,a,\boldsymbol{\theta})$.

#### Application

The data $\mathcal{D}$ were generated from the Gaussian distributions in Eqs (1-4). The following specifications of parameters were used:

$$
n_0 = n_1 = 500, \quad \mu_1 = 12, \quad \mu_0 = 7, \quad \sigma_1^2 = 4, \quad \sigma_0^2 = 4,
$$

$$
\rho_0 = -0.7, \quad \rho_1 = -0.7, \quad \mu_{a1} = 15, \quad \mu_{a0} = 10, \quad \sigma_{a1}^2 = 9, \quad \sigma_{a0}^2 = 4,
$$

$$
\beta_0 = \rho_0\frac{\sigma_{a0}}{\sigma_0}, \quad \alpha_0 = \mu_{a0} - \beta_0 b_0, \quad \beta_1 = \rho_1\frac{\sigma_{a1}}{\sigma_1}, \quad \alpha_1 = \mu_{a1} - \beta_1 b_1,
$$

$$
\sigma_3^2 = \sigma_{a0}^2(1-\rho_0^2), \quad \sigma_4^2 = \sigma_{a1}^2(1-\rho_1^2).
$$

Here the auxiliary parameters, $\mu_{a0}, \mu_{a1}, \sigma_{a0}^2, \sigma_{a1}^2$, give the marginal means and variances for marginal means and variances of $A_0, A_1$, and $\rho_i$ is the correlation between $A_1$ and $B_i$, $(i = 0, 1)$. These parameters were introduced so that the simulated data are generated from a bivariate Gaussian distribution, but they play no direct part in the generative models of $A_i$ given $B_i = b_i$ and $Z_2 = i, (i = 0, 1)$.

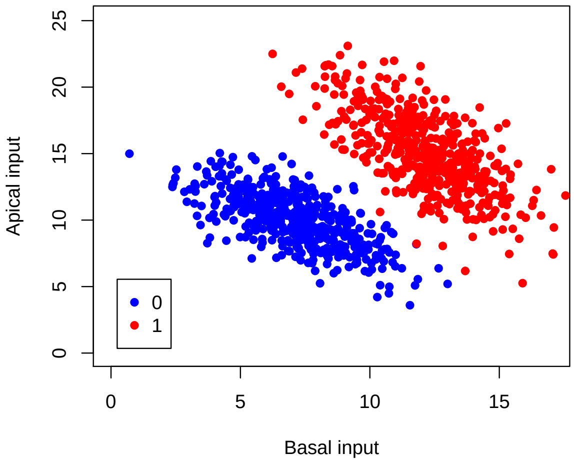


Figure 1: Simulated basal and apical data. The points in red are those where $Z_2 = 1$, while the points in blue are those where $Z_2 = 0$.

The simulated data are displayed in Figure 1.

By using the procedures for simulating from $p(\boldsymbol{\theta}|\mathcal{D})$ in Section 5, we can generate $N$ random draws and use the equations in Section 6 to find estimated posterior probabilities of a second AP for new values $b, a$, given the data $\mathcal{D}$, and also to compute credible intervals for the posterior probability of a second AP.

An R script Supp-S2.R is available to perform the computation. It is available from the repository

https://github.com/JWKay/Pyramid

This script was used to generate the data for the following contour and surface plots of the estimated posterior probability of a second AP for a grid of values of basal and apical input. The plots were produced using Mathematica. The units are arbitrary.

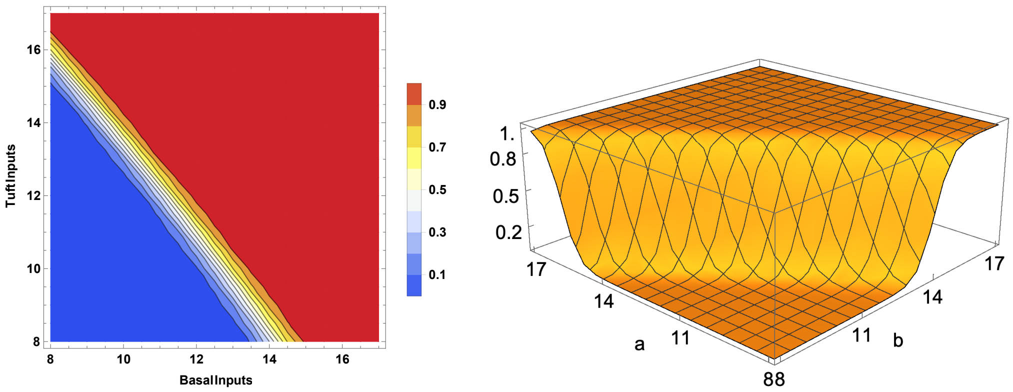

#### Some estimated posterior probabilities and credible intervals

The following table gives some particular estimates of posterior probability for given basal and apical inputs, as well as estimated Monte Carlo standard errors, and 0.05th , 0.5th and 0.95th quantiles of the simulated posterior distribution of $P(S_2|b, a, \boldsymbol{\theta})$.

| | $b$ | $a$ | $\hat{P}(S_2\|b, \mathcal{D})$ | $\hat{P}(S_2\|b, a, \mathcal{D})$ | MCSE | $q_{0.05}$ | $q_{0.5}$ | $q_{0.95}$ |
|---|---|---|---|---|---|---|---|---|
| Case 1 | 10 | 13 | $8.48 \times 10^{-3}$ | 0.198 | $7.40 \times 10^{-4}$ | 0.0974 | 0.186 | 0.206 |
| Case 2 | 12 | 11 | 0.118 | 0.536 | $1.20 \times 10^{-3}$ | 0.339 | 0.535 | 0.734 |
| Case 3 | 14 | 10 | 0.653 | 0.987 | $1.09 \times 10^{-4}$ | 0.966 | 0.990 | 0.997 |

A credible interval is useful for assessing the uncertainty in an estimate $\hat{P}(S_2|b, a, \mathcal{D})$ of the posterior probability $P(S_2|b, a, \mathcal{D})$. From the table, we see that for Case 1 the estimated posterior probability of a second AP is 0.198, and the 90% credible interval is (0.0974, 0.206). Thus we can be at least 90% certain that $P(S_2|10, 13, \mathcal{D})$ is less than 0.206. For Case 2, there is much uncertainty with an estimated posterior probability of 0.536, and a very wide credible interval. In Case 3, we can be at least 90% certain that the posterior probability $P(S_2|14, 10, \mathcal{D})$ is greater than 0.966.

#### Application of the Bayesian interpretation

We now discuss the Bayesian interpretation in terms of posterior odds, by making use of the estimated posterior probabilities. The prior probability of a second AP is assumed to be 0.005, so the prior odds in favour of $S_2$ are $1/199 \doteq 0.00503$, with log odds approximately -5.2933. For Case 1, taking the basal input into account gives the posterior odds in favour of $S_2$ as 0.00856, while the posterior odds given both the basal and apical input are increased to 0.246. The corresponding posterior odds for Case 2 are from 0.00503 to 0.0.134 to 1.15. For Case 3, the odds change from 0.00503 to 1.88, given the basal input of 14, and then increase to 74.3, given also the apical input of 10.

#### Better generative models?

This illustration has employed simple models. Nevertheless, the contour and surface plots above seem fairly plausible as an approximation. It seems likely, however, that the boundary between the 0s and 1s could be nonlinear.

On the range of values of basal input, say $[0, M_b]$, one could expect that the conditional distribution of basal input given that $Z_2 = 0$ would peak at 0 and be skewed to the right, and that the conditional distribution of basal input given that $Z_2 = 1$ would peak at $M_b$ and be skewed to the left. Suitable truncated Gaussian probability distributions could be used as generative models for the basal input to reflect these assumptions.

In the generative models for the apical input, on a grid $[0, M_a]$ say, given the basal input and the value of $Z_2$, one could expect the following. Given $Z_2 = 0$, the conditional mean of apical input is likely to decrease in a nonlinear manner from a value that is the marginal mean of the apical input towards zero, possibly along a four-parameter logistic curve, with two asymptotes. Given $Z_2 = 1$, the conditional mean of apical input is likely to decrease in a nonlinear manner from a value that is, say, the 0.95 quantile of the apical input towards the marginal mean of the apical input, possibly along a different four-parameter logistic curve, also with two asymptotes. Given $Z_2 = 0$, when the basal input is very low one might expect the spread of the apical values to be large and then become smaller as the level of basal input increases. Therefore, the conditional variance of apical input given $Z_2 = 0$ could be a decreasing function of basal input. Given $Z_2 = 1$, when the basal input is very low one might expect the spread of the apical values to be small and then become larger as the level of basal increases. Therefore, the conditional variance of apical input given $Z_2 = 1$ could be an increasing function of basal input. Therefore, the generative models for the apical input given the basal input could be weighted nonlinear regression models, with weights a function of basal input, and with Gaussian errors.
