## Supplementary File S3 for "Bayesian modeling of BAC firing as a mechanism for apical amplification in neocortical pyramidal neurons"

### Activation Functions

#### Derivation

Consider a local processor which possesses two sites of integration. One site receives input from a receptive field, while the other receives input from a contextual field. Let the integrated weighted and summed inputs be $r, c$, respectively. Such processors were used in artificial neural nets in the 1990s.

In order to produce an output these two integrated fields must be combined, but how? The following four requirements were set and the aim then was to invent a function A(r, c) which could satisfy all four properties.

1. If the integrated RF input is zero then the activation is zero.
2. If the integrated CF input is zero then the actvivation should be the integrated RF input.
3. If the integrated RF and CF inputs have the same sign, then the activation should be greater than when it is based on the RF input alone. On the other hand, if the integrated RF and CF inputs disagree, i.e. are of opposite sign, then the activation should be less than it would be if based on the RF input alone.
4. The sign of the activation should be that of the integrated RF so that the context cannot affect the direction of the output decision.

Requirements 1 says that $A(0, c) = 0$, while requirement 2 is that $A(r, 0) = r$.

These requirements, taken together with requirement 4, suggest that $A(r, c)$ should have the form $rA1(r, c)$, where $A1$ is a positive function with $A1(r, 0) = 1$.

Requirements 3 & 4 suggest now that $A1(r, c)$ should have the form $A2(rc)$, where $A2$ is a function of a single variable and is greater than unity when its argument is positive and less than unity when its argument is negative, and also $A2(0) = 1$. The exponential function has these properties, and so we now consider the function

$$A(r, c) = r \exp(mrc)$$

where $m > 0$.

This form for $A(r, c)$ has the property that when $rc$ is very negative it approaches 0. This means that in cases of very strong disagreement between the integrated RF and CF inputs the output is then placed in its most uncertain state. However, because we wish the RF inputs to be primary drivers, we don't necessarily desire that the RF information in such a combination is entirely ignored. Hence we introduce a constant $s$ which plays the role of a depression factor; that is, in the extreme case when $rc \longrightarrow -\infty$, the activation will be $sr$. These requirements lead naturally to the following class of activation functions

$$A(r,c) = r[s + (1-s)\exp(mrc)] = sr + (1-s)r\exp(mrc) = r + (1-s)r[e^{mrc} - 1],$$

with $m > 0$ and $0 \le s < 1$. This suggests the use of the general activation functions

$$f(x) = x, \quad \text{and} \quad g(x,y) = (1-s)x[e^{mxy} - 1],$$

These activations functions are used with various choices of $x, y, s, m$ in the manuscript. We now illustrate the use of these functions by formulating the general activation function used in the general model.

#### Application: activation function for the general model

Here, $a$ and $b$ denote new values of the apical and basal input, respectively. In the apical site, we take $x = a, y = z_1, s = \frac{1}{2}, m = \beta_4$ so that the activation is

$$\tfrac{1}{2}a(1 + \exp{(\beta_4 a z_1)}).$$

When an initiating backpropagated AP arrives in the apical site then $z_1 = 1$ and

$$c = \tfrac{1}{2}a(1 + \exp{(\beta_4 a)}).$$

When BAC firing has been triggered, this value of $c$ arrives in the somatic site. Then, with $x = b, y = c, s = \frac{1}{2}, m = \beta_3'$, the somatic activation function is

$$\tfrac{1}{2}b[1 + \exp{(\beta_3' bc)}].$$

Thus, the expression for the activation function becomes

$$\tfrac{1}{2}b[1 + \exp[\tfrac{1}{2}\beta_3' ba(1 + \exp{(\beta_4 a)})]].$$

We then take the general activation functions to be

$$\beta_2' * \tfrac{1}{2}b[1 + \exp[\tfrac{1}{2}\beta_3' ba(1 + \exp{(\beta_4 a)})]] \equiv \beta_2 b[1 + \exp[\beta_3 ba(1 + \exp{(\beta_4 a)})]],$$

where $\beta_2 \equiv \frac{1}{2}\beta_2'$ and $\beta_3 \equiv \frac{1}{2}\beta_3'$. This activation function is used in the general model that is fitted to the binarised data from Shai et al. [10].
