## Supplementary figures and images for "Bayesian modeling of BAC firing as a mechanism for apical amplification in neocortical pyramidal neurons"

### Supplementary Figure S1

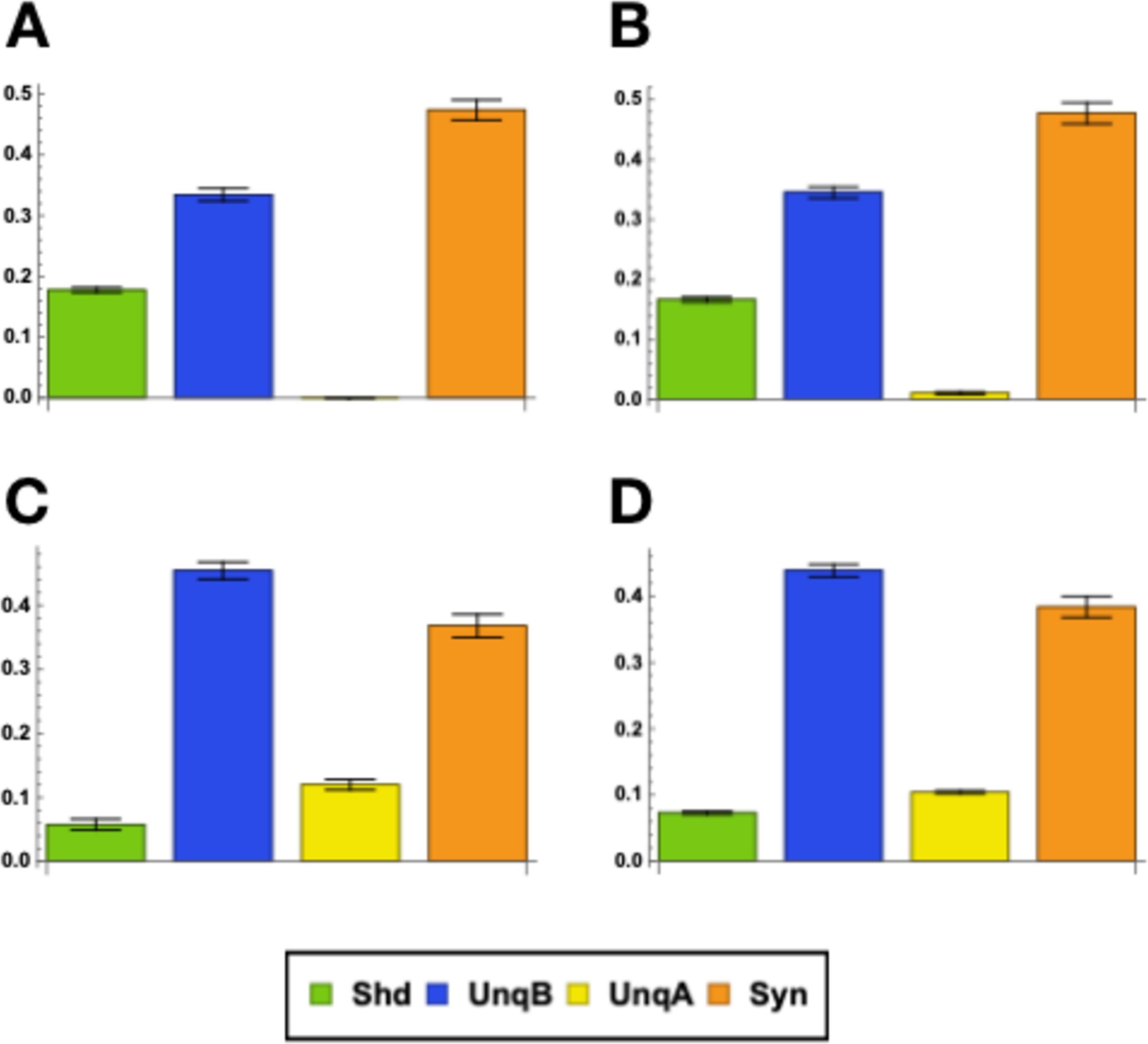
